# Inhibition of the SHP-1 activity by PKC-θ regulates NK cell activation threshold and cytotoxicity

**DOI:** 10.1101/2021.09.06.459131

**Authors:** Aviad Ben-Shmuel, Batel Sabag, Guy Biber, Abhishek Puthenveetil, Moria Levy, Tammir Jubany, Noah Joseph, Omri Matalon, Jessica Kivelevitz, Mira Barda-Saad

## Abstract

Natural Killer (NK) cells play a crucial role in immunity, killing virally infected and cancerous cells. The balance of signals initiated upon engagement of activating and inhibitory NK receptors with cognate ligands determines killing or tolerance. Nevertheless, the molecular mechanisms regulating rapid NK cell discrimination between healthy and malignant cells in a heterogeneous tissue environment are incompletely understood. The SHP-1 tyrosine phosphatase is the central negative NK cell regulator, which dephosphorylates key activating signaling proteins. Though the mechanism by which SHP-1 mediates NK cell inhibition has been partially elucidated, the pathways by which SHP-1 is itself regulated remain unclear. Here, we show that phosphorylation of SHP-1 in NK cells on the S591 residue by PKC-θ promotes the inhibited SHP-1 “folded” state. Silencing PKC-θ maintains SHP-1 in the active conformation, reduces NK cell activation and cytotoxicity, and promotes tumor progression *in-vivo*. This study reveals a molecular pathway that sustains the NK cell activation threshold through suppression of SHP-1 activity.

## Introduction

Natural Killer (NK) cells are a class of lymphocytes that comprise a central component of the innate immune system. They patrol the body and kill virally transformed and cancerous cells. NK cell effector functions are carried out by the release of cytotoxic granules containing perforin and Granzyme-B, which induce target cell death(Topham *et al*., 2009),(Pipkin *et al*., no date). Furthermore, NK cells produce cytokines and chemokines, such as IFN-γ, which activate the adaptive immune response(Mαληοτρα *ετ αλ.*, 2011).

Activation or inhibition of NK function was shown to depend on the balance between signals derived from activating and inhibitory receptors on the NK cell surface(Lanier, 2005). Various motifs on the cytoplasmic tails of these receptors were shown to induce distinct downstream signaling cascades. These include immunoreceptor tyrosine-based activation motifs (ITAMs) found on cytoplasmic tails of the transmembrane adaptor proteins CD3ζ, FcεRIγ, and DAP12, or the YINM motif on the DAP10 adaptor, which associates with the NKG2D receptor(André *et al*., 2004). Another group of motifs includes the immunoreceptor tyrosine-based switch motifs (ITSMs) on the CD244 receptor(Lanier, 2003),(Feng *et al*., 2005),(Vyas *et al*., 2002b). Though NK cell receptors are well characterized, the molecular mechanisms downstream to receptor ligation, and how these are integrated to achieve the appropriate activation state, are incompletely understood.

Inhibitory NK cell receptors express a separate target sequence for signal suppression, consisting of immunoreceptor tyrosine-based inhibition motifs (ITIMs) expressed on the cytoplasmic tails(Long, 2008). The key receptors facilitating NK inhibition recognize Major Histocompatibility Class 1 (MHC-I) molecules expressed on the majority of cells in the body. These receptors include the Killer-cell Immunoglobulin-like Receptor (KIR) family in humans and primates, the Ly49 family in mice, and NKG2A/CD94 receptors expressed in both primates and mice(Orr *et al*., 2010). NGK2A/CD94 were shown to have specificity for the Human Leukocyte Antigen (HLA) isoform E (HLA-E), whereas the KIR family of inhibitory receptors was shown to associate with HLA-C and HLA-B(Moretta *et al*., 1994),(Wagtmann *et al*., 1995). Engagement of these receptors with their cognate ligands and phosphorylation of their ITIM motifs recruits the Src-homology 2-domain (SH2)-containing protein tyrosine phosphatase (PTP), SHP-1. SHP-1 consequently dephosphorylates key proteins involved in the activating signaling pathway, thereby downregulating NK cell activation(Stebbins *et al*., 2003; Matalon *et al*., 2016). SHP-1 is comprised of a catalytic phosphatase domain near its C’ terminus, two phospho-tyrosine binding SH2 domains towards its N’ terminus, and sites of tyrosine and serine phosphorylation at the end of its C’-terminal tail(Lorenz, 2009). In its auto-inhibited closed state, the SHP-1 N’ SH2 domain associates with the catalytic domain, blocking access to its substrates.

Binding of SHP-1 SH2 domains to phosphorylated tyrosine residues on the ITIM of the inhibitory receptors induces a conformational alteration in SHP-1. This releases it from the inhibited state, and enables its catalytic activity, i.e. dephosphorylation of key signaling molecules and inhibition of NK cell function(Wang *et al*., 2011). SHP-1 substrates in NK cells include the guanine nucleotide exchange factor (GEF), VAV1(Stebbins, 2003), Phospholipase C gamma 1/2 (PLCγ1/2), and Linker for Activation of T-cells (LAT), recently identified by our group(Matalon, 2016).

Though SHP-1 function is critical for correct NK cell immune responses, the manner by which its catalytic activity is regulated remains unclear. Elucidating this mechanism is especially important, since NK cells engage target cells expressing both activating and inhibitory receptors, and SHP-1 recruitment to ITIMs decreases the activity of NK cell towards potential targets, despite expression of activating ligands. Therefore, upregulation of ligands for inhibitory NK receptors that express ITIMs by cancer cells, such as HLA-E and PD-L1, may repress NK cell activity through SHP-1 recruitment and activity(Carretero *et al*., 1998; Chemnitz *et al*., 2004). Identifying the factors that regulate SHP-1 activity may lead to a better understanding of how NK cell activity is accurately and rapidly tuned in tissues containing both healthy and transformed cells. We recently revealed that actin retrograde flow (ARF) induces a physical conformational change in the SHP-1 molecule to facilitate its activity(Matalon *et al*., 2018). However, key questions remain unresolved. SHP-1 was shown to localize to the lytic NK cell synapse (NKIS)(Vyas *et al*., 2002a), yet the mechanism by which its activity within the lytic synapse is regulated was not determined. Furthermore, it is not clear how termination of SHP-1 activity occurs after NK cells are successfully inhibited, thereby enabling NK cells to function in subsequent interactions with activating target cells.

Thus, since SHP-1 is a critical checkpoint molecule in NK cells and regulates the NK cell activation threshold, it is of great interest to understand the underlying mechanisms that control its catalytic activity during the NK cell response.

Here, we demonstrate that in NK cells, SHP-1 is heavily phosphorylated on the serine 591 residue (S591) via PKC-θ (and hence dormant) during activating but not inhibitory responses, within the first 5 minutes of NK cell activation. This phosphorylation is dynamic, and increases along the progression of the inhibitory NKIS, while slightly decreasing during progression of the activating NKIS. SHP-1 phosphorylation dynamics were also closely correlated with changes in SHP-1 catalytic conformations. Blocking PKC−θ-mediated phosphorylation of SHP-1 restored SHP-1 activity and inhibited NK cell activation. Furthermore, NK cells bearing a phosphor-mimetic point mutation that suppresses SHP-1 activity 1 at the serine 591 residue were highly activated. Finally, knockout of SHP-1 in NK cells rescued PKC-θ silencing and promoted tumor clearance *in-vivo*.

The PKC-θ-mediated regulation of SHP-1 potentially serves to maintain and prime NK cellsin a complex cellular environment consisting of both healthy and malignant cells, requiring finely tuned localized NK cell activity.

## Results

### SHP-1 S591 phosphorylation in NK cells is a dynamic process, differentially regulated during inhibitory and activating interactions

It remains unclear how SHP-1 activity is regulated throughout the duration of the NK cell response, and how this regulation is maintained in different states, i.e. during NK cell activation and inhibition. Furthermore, SHP-1 is recruited to both cytolytic as well as non-cytolytic NK synapses, demonstrating that different modes of regulation are needed to ensure proper NK cell responses(Vyas *et al*., 2001; Vyas, 2002a). Phosphorylation of SHP-1 was previously shown in different cellular systems(Li *et al*., 1995; Jones *et al*., 2004; Liu *et al*., 2007). The outcome of these molecular processes, however, demonstrated contradictory results, and molecular regulation of SHP-1 in NK cells has not been fully addressed. We recently employed a mutant YTS-2DL1 knock-in line expressing the SHP-1 phosphor-mimetic serine to aspartic acid residue substitution (SHP-1 S591D), which exhibits increased anti-tumor NK function relative to WT SHP-1-expressing YTS cells(Ben-Shmuel *et al*., 2020). To dissect the possible effect of SHP-1 phosphorylation on NK cell function, physiological activating and inhibiting interactions were induced with 721.221 target cells, and S591 phosphorylation patterns were examined in YTS-2DL1 cells and isolated primary NK KIR2DL1^+^ cells (pNK-2DL1) from healthy human donors. We conducted functional assays as previously described(Matalon, 2016, 2018) by incubating YTS-2DL1 or primary NK cells (pNKs cells) isolated from PBMCs of healthy donors expressing KIR2DL1+ (referred to as pNK KIR2DL1) with 721.221 target cells either expressing the KIR2DL1 cognate ligand, HLA Cw4 (721-Cw4, which inhibits NK activity), or an irrelevant HLA Cw7 ligand (721-Cw7 or 721-HLA negative cells(Münz *et al*., 1997), which promote NK cell activation). Cell lysates were immunoblotted with anti-pSHP-1 S591 antibody. Our data revealed different SHP-1 phosphorylation profiles during NK cell inhibition and activation after 5 minutes of incubation (Fig. 1A). High S591 phosphorylation levels could be seen during activating interactions (3.06±0.31*,P*=0.0104), whereas lower SHP-1 S591 phosphorylation was observed during induced inhibitory interactions. The same pattern was observed during incubation of pNK-2DL1 cells with activating 721-HLA negative cells or with inhibiting 721-Cw4 targets cells (2.51±0.19*,P*=0.006, Fig. 1B). The formation of the immunological synapse (IS) is highly dynamic, involving movement, activation, and termination of signaling complexes(Burroughs *et al*., 2002). Therefore, we wished to analyze the change in SHP-1 S591 phosphorylation over time. YTS-2DL1 or pNK-2DL1 cells were incubated with activating or inhibiting 721.221 targets for 5, 10, 15 and 20 minutes. Strikingly, we found that pS591 on SHP-1 was dramatically altered <u>during formation</u> of the inhibitory NK cell synapse, showing almost no initial phosphorylation after 5 minutes of incubation, and displaying high phosphorylation by 20 minutes. Activating NK cell interactions, however, displayed higher SHP-1 S591 phosphorylation during the first 5 minutes of activation, remaining relatively stable, with a slight (non-significant) reduction after 20 minutes of activation (Fig. 1C and Fig 1-Fig. supplement 1). Collectively these results suggest that SHP-1 S591 phosphorylation may play a role during NK cell activation, and during late inhibitory interactions. This mechanism may attenuate SHP-1 functionality in order to enable NK cell activation within these time frames.

**Fig. 1.**
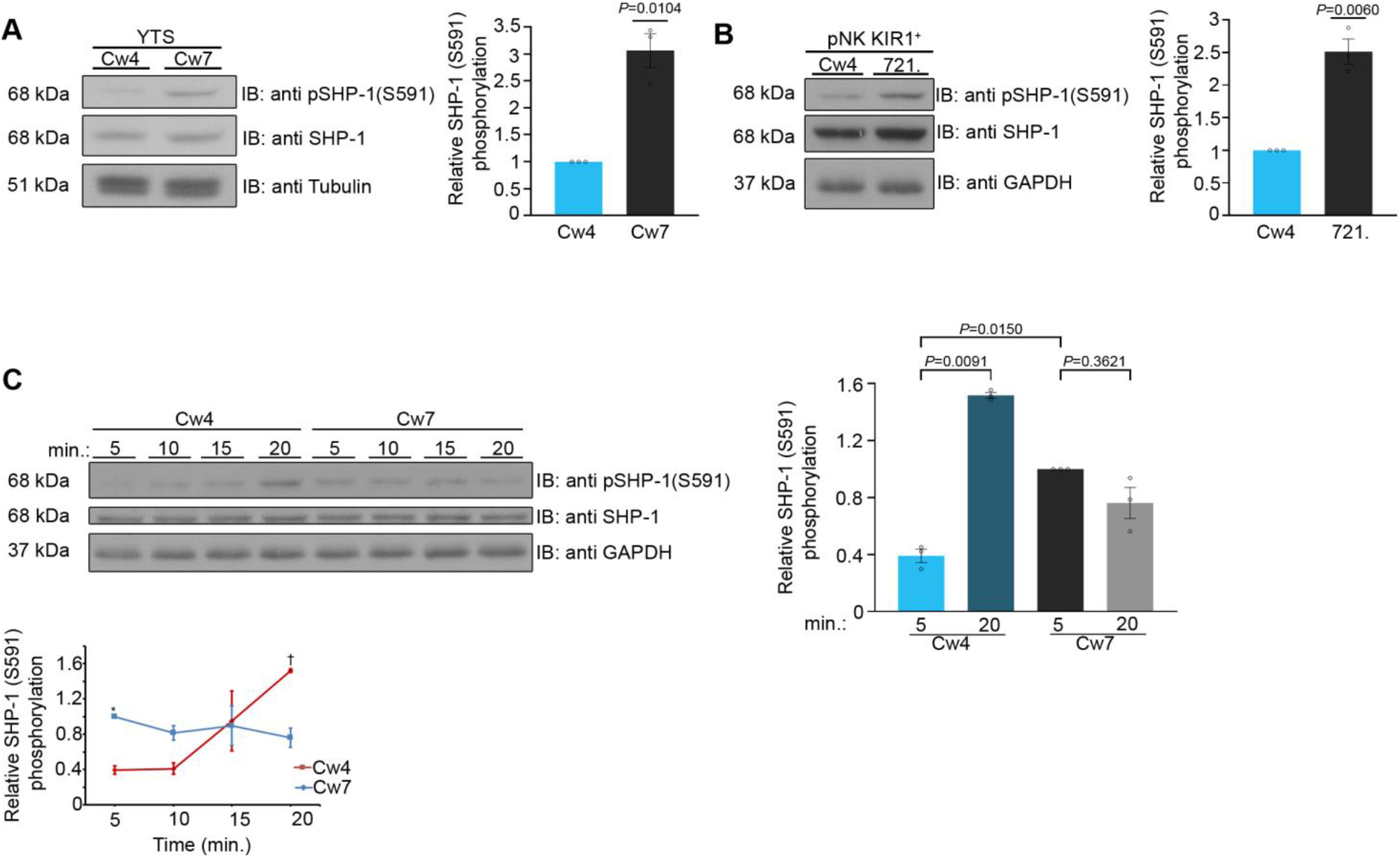
Phosphorylation kinetics of SHP-1 S591 during activating and inhibitory NK cell interactions. **(A)** YTS-2DL1 NK cells were incubated with either inhibitory 721-Cw4 HLA or 721-Cw7 HLA target cells at 37°C for 5 minutes, and then lysed. Lysates were separated on SDS page and immunoblotted with anti-pSHP-1 S591 antibody. SHP-1 S591 phosphorylation levels were measured by densitometric analysis, relative to -Tubulin loading control using ImageJ. Samples were normalized to the YTS-2DL1 sample incubated with 721-Cw4 target after 5 minutes of activation (*P*=0.0104, quantification on the right showing the average of 3 independent experiments). **(B)** pNK-2DL1 cells were incubated with either 721-Cw4 HLA or 721-HLA negative target cells at 37°C for 5 minutes. pSHP-1 S591 levels were determined as in (a) (*P*=0.0060, quantification on the right showing the average of 3 independent experiments). **(C)** YTS-2DL1 cells were incubated with target cells as described in (a), for four different time points, as indicated. pSHP-1 S591 levels were quantitated as in (a). Statistical significance between Cw4 and Cw7 after 5 minutes of activation (*P*=0.015), statistical significance between Cw4 at 5 minutes versus 20 minutes (*P*=0.0091). pSHP-1 S591 levels of YTS-2DL1 cells incubated with targets for 5 and 20 minutes are shown in the bar graph (quantification showing the average of 3 independent experiments). Data are shown as mean ± SEM. One sample t-tests (A, B) or one-way ANOVA with Tukey test (C) were used to calculate p values **Figure 1-source data 1: Representative blots** **Figure 1-source data 2 and and Figure 1—figure supplement 1-source data 1: Numerical data for all graphical presentation.**

### SHP-1 conformational kinetics parallel S591 phosphorylation patterns

To elucidate whether SHP-1 phosphorylation on the S591 residue complements SHP-1 conformation and activation status, a SHP-1 Förster resonance energy transfer (FRET) bio-sensor construct was cloned into YTS-2DL1 cells as we previously described(Matalon, 2018). The FRET sensor was constructed with SHP-1 tagged on the N’ and C’ termini with YFP and CFP, respectively (YFP-SHP-1-CFP). It is known that SHP-1 activation status is correlated with its conformation(Poole *et al*., 2005; Wang, 2011). In the closed conformation the N’ SH2 domain masks the catalytic domain rendering the enzyme inactive, whereas when the catalytic domain is free of the N’ SH2 domain, the protein remains in an open active conformation(Wang, 2011). Hence, an inactive SHP-1 protein provides high FRET efficiency due to YFP and CFP proximity, and an active SHP-1 protein demonstrates low FRET efficiency as SHP-1 is in an open conformation, distancing the two reporter proteins. With this construct, we could image the dynamic activation and inhibition status of SHP-1 throughout the lifetime of the NKIS as previously reported(Matalon, 2018). YTS-2DL1 cells were transfected with the YFP-SHP-1-CFP FRET sensor construct and incubated with inhibiting 721-Cw4 or activating 721-Cw7 cells stably expressing mCherry in order to distinguish NK and target cells. FRET efficiency was measured using high resolution microscopy, as previously described(Barda-Saad *et al*., 2005; Pauker *et al*., 2012; Fried *et al*., 2014). In this set of experiments, we chose to focus on very early (5 minutes) and late (20 minutes) stages of NK cell: target cell interaction, as they showed the largest change in SHP-1 S591 phosphorylation (Fig. 1 and Fig. 1-Fig. supplement 1). FRET measurements demonstrated patterns similar to SHP-1 S591 phosphorylation profiles; during initial inhibitory NK:721-Cw4 interactions (5 minutes), SHP-1 in NK cells displayed low synaptic FRET efficiency, indicating an open and active state (8.62%±1.8), shifting to high FRET efficiency after 20 minutes of synapse maturation (15.13%±1.9, *P*=0.0300; Fig. 2A&B top two panels), indicating a closed and inhibited state. During activating interactions, however, SHP-1 in NK cells displayed highest synaptic FRET efficiency after 5 minutes (18.45%±1.9, *P*=0.0003 comparing 5 minutes of activation to inhibition), indicating a closed and inhibited state, while after 20 minutes of activation, a decrease was observed in SHP-1 FRET efficiency to similar levels as NK:721-Cw4 interactions after 5 minutes (8.15%±1.9, *P*=0.006; Fig. 2A&B bottom two panels), indicating an open and active SHP-1 conformation(Matalon, 2018). These data demonstrate that SHP-1 conformation, and accordingly, its activation status, changes differentially during activating and inhibiting interactions as the NKIS matures, correlating with S591 phosphorylation patterns. Our results suggest that SHP-1 conformation and catalytic activation are regulated in a temporal manner at both the activating and inhibitory NKIS, and suggest a possible role for SHP-1 S591 phosphorylation on SHP-1 activity in NK cells.

**Fig. 2.**
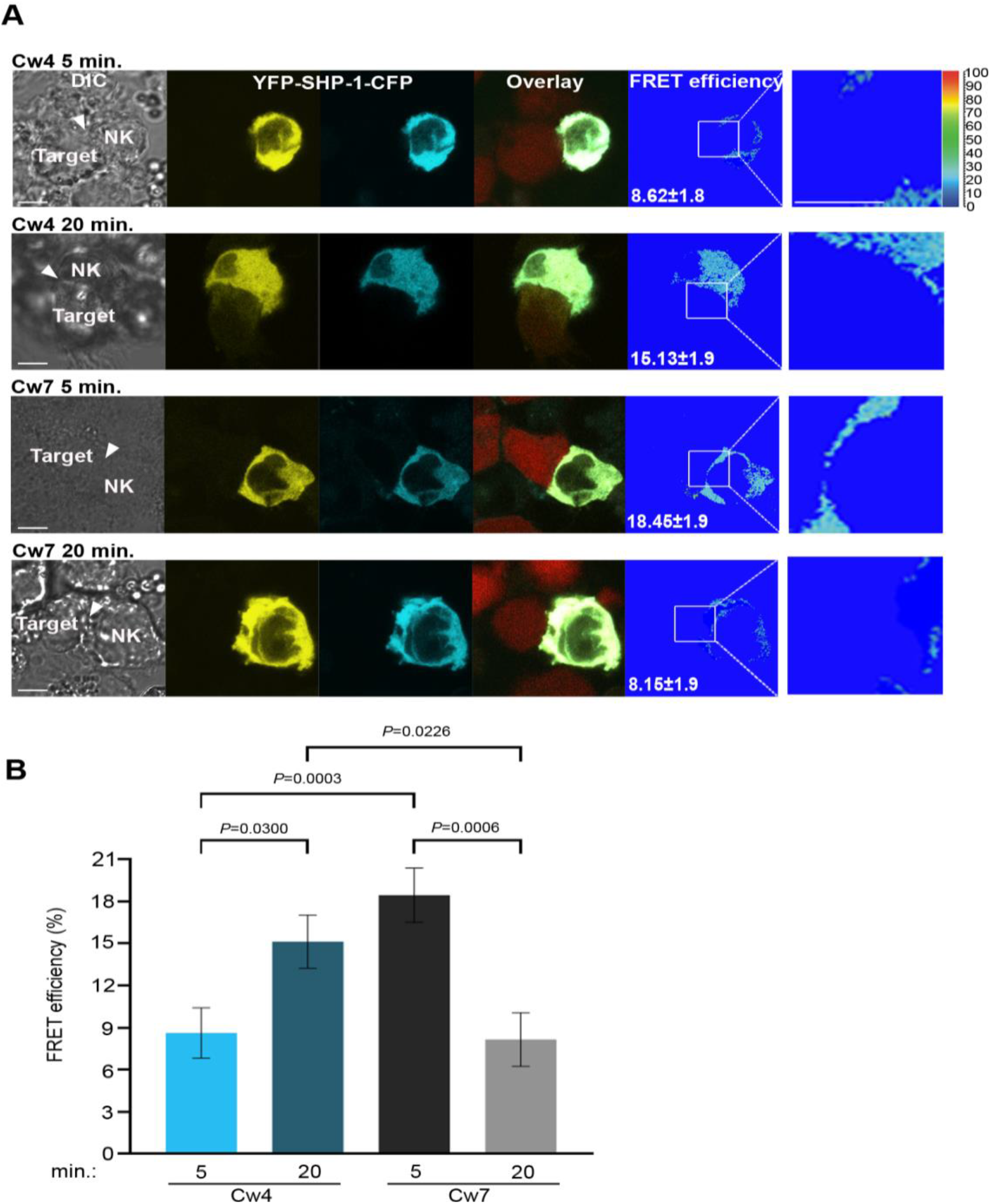
SHP-1 conformational dynamics reflect S591 phosphorylation during activating and inhibitory NK cell interactions. **(A)** YTS-2DL1 YFP-SHP-1-CFP cells were incubated over slides pre-seeded with 721-Cw4 (top panels) or Cw7 (bottom panels) target cells expressing mCherry. The cells were incubated for 5 or 20 minutes at 37°C to enable conjugate formation, and fixed. FRET analysis was performed as indicated. **(B)** Graph summarizing FRET efficiency following 5 or 20 minute activation with Cw4 or Cw7 target cells. For Cw4, 5 and 20 minutes activation, n=72 and 62 cell conjugates analyzed, respectively. For Cw7, 5 and 20 minutes activation, n= 73 and 47 cell conjugates analyzed from three independent experiments, respectively. Data are shown as mean ± SEM. Two-way ANOVA with Tukey test (B) were used to calculate p values **Figure 2-source data 1: Numerical data for the graphical presentation.**

### pSHP-1 distinguishes between NK cell activating and inhibiting synapses, and correlates with PKC-θ accumulation at the NKIS

In transformed tissues, NK cells interact with both susceptible targets and bystander cells which may both influence NK cell function(Zhou *et al*., 2017). NK cells are capable of serial cytotoxicity, and rapidly engage with and detach from these different targets(Choi *et al*., 2013; Forslund *et al*., 2015; Guldevall *et al*., 2016). It was recently demonstrated that NKIS maintenance is highly regulated, and signaling events which lead to NK cell attachment to new target cells accelerate the detachment from previous targets(Netter *et al*., 2017; Srpan *et al*., 2018). Furthermore, highly regulated lytic granule convergence to the MTOC is crucial for NK cells to avoid bystander cell killing(Hsu *et al*., 2016). Thus, it is evident that controlled and coordinated signaling events are critical for NK cell target identification and subsequent precise function.

We hypothesized that SHP-1 S591 phosphorylation may thus enable rapid discrimination by NK cells between healthy and malignant targets, facilitating controlled and sequential killing in a heterogeneous environment. To examine whether SHP-1 phosphorylation in NK cells is coordinately directed when NK cells are challenged simultaneously with activating and inhibitory signals, pNK-2DL1 cells were concurrently incubated with both activating K562-CFP and inhibiting 721-Cw4-mCherry stably labelled cells. Synapse intensity of pSHP-1 S591 was assessed in NK cells that were found forming simultaneous dual synapses with both activating and inhibiting cells. Strikingly, NK cells were able to rapidly distinguish between activating and inhibitory targets cells as measured by higher pSHP-1 S591 levels that were localized to synapses with K562, versus 721-Cw4 cells (Fig. 3A, *P*=0.0074). Hence SHP-1 S591 phosphorylation is a regulated and directed event and may allow NK cells to locally regulate SHP-1 phosphorylation to enable the maintenance of multiple local activation states in a single cell.

**Fig. 3.**
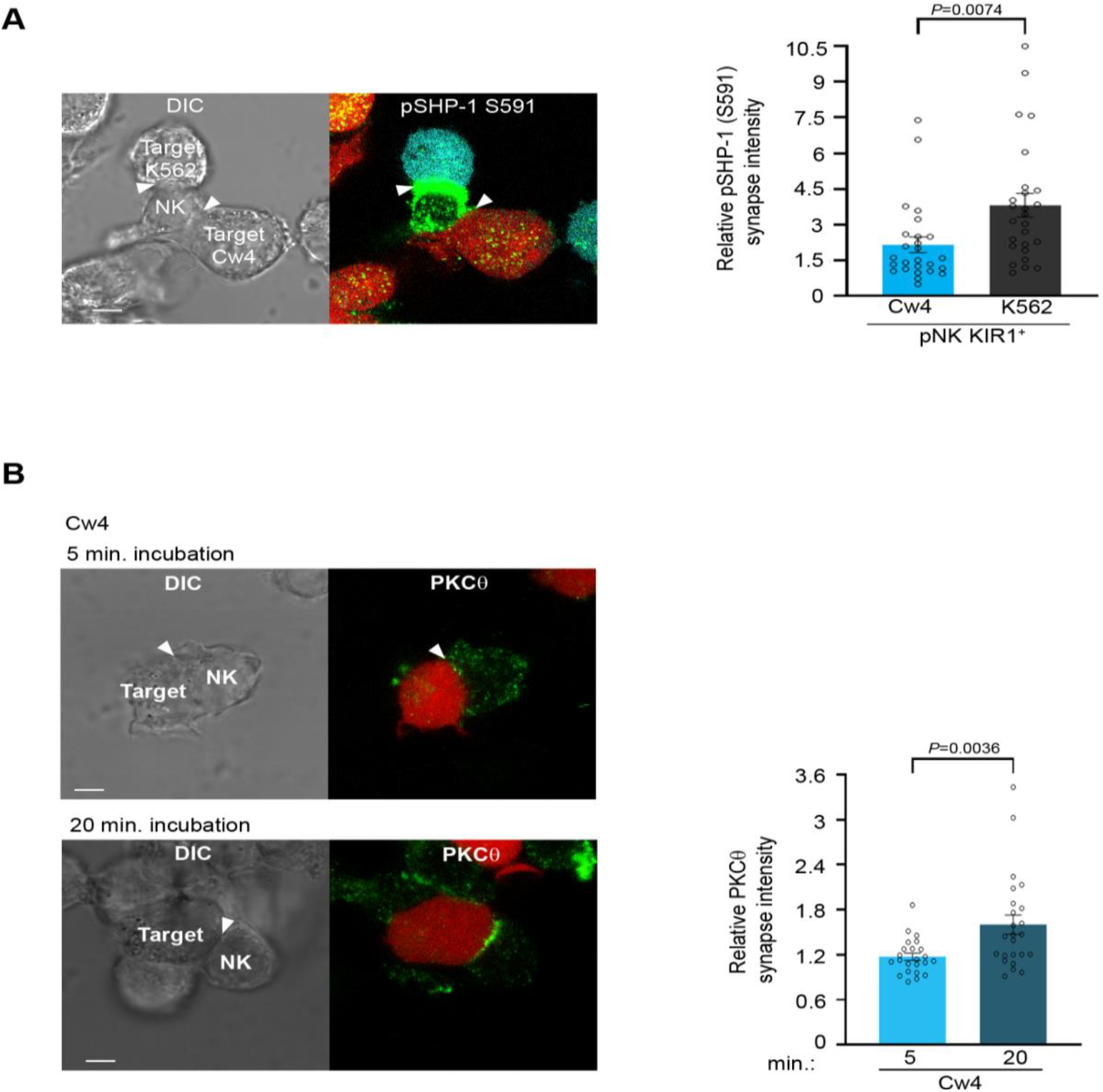
Phosphorylation of SHP-1 S591 occurs at the activating NKIS during simultaneous activating and inhibitory synapse formation, in parallel to PKC-θ accumulation. **(A)** pNK-2DL1 cells were incubated over slides pre-seeded with 721-Cw4 and K562 cells expressing mCherry or CFP, respectively. The cells were incubated for 5 minutes at 37°C to enable conjugate formation, and were fixed. pSHP-1 S591 was labeled with primary rabbit anti pSHP-1 S591 antibody, and secondary anti-rabbit 488 antibody. Synapse intensity was quantified in NK cells relative to each cell in multiple NK cell synapses with two different targets (*P*=0.007, quantification on the right of triple cell conjugates collected, n=26). **(B)** YTS-2DL1 cells were incubated over slides pre-seeded with 721-Cw4 target cells expressing mCherry. The cells were incubated for 5 or 20 minutes at 37°C to enable conjugate formation, and were fixed. PKC-θ was subsequently labeled with primary goat anti PKC-θ antibody, and secondary anti-goat 488 antibody. **(C)** Graph summarizing PKC-θ accumulation at the NKIS at two activation times. Analysis was conducted comparing PKC-θ intensity at the NKIS relative to the rest of the NK cell. For Cw4 5 and 20 minute activation, n=24 cell conjugates were analyzed from three independent experiments. Data are shown as mean ± SEM. One sample t-tests (A, B) were used to calculate p values **Figure 3-source data 1 and Figure 3-Figure supplement 1-source data 2: Numerical data for all graphical presentation.** **Figure 3-Figure supplement 1-source data 1: Representative blots**

We next aimed to identify the serine kinase implicated in SHP-1 S591 phosphorylation. PKC-θ is expressed predominantly in hematopoietic and muscle cells(Osada *et al*., 1992; Baier *et al*., 1993), and was shown to play multiple roles in T-cell activation(Hayashi *et al*., 2007). In human NK cells, the role and molecular pathways of PKC−θ are incompletely defined. PKC−θ was suggested to participate in murine NK cell activity(Tassi *et al*., 2008; Aguiló *et al*., 2009; Anel, Juan I Aguiló, *et al*., 2012; Merino *et al*., 2012), however the mechanism by which it exerts this function remains unclear. It was shown that TNF-α and IFN-γ secretion is defective in PKC-θ^−/−^ mice(Page *et al*., 2008), and this can contribute to defective recruitment of effector cells to the site of tumor development. In a different study(Tassi, 2008), no effect of PKC-θ deficiency was observed on IFN-γ secretion induced by IL-12, IL-18, or a combination of both cytokines. Therefore, it is not clear how PKC-θ is involved in the regulation of NK cell signaling cascades, and if its activity impacts human NK cell function.

Early reports by Vyas et al showed that PKC-θ localizes to the cytolytic NK synapse during early activation, i.e. following 5 and 10 minutes of activating target cell conjugation, and relatively low PKC-θ polarization is observed following 10 minutes of non-cytolytic NK: target cell conjugates(Vyas, 2002a). This PKC-θ localization was similar to the localization of SHP-1 at the NKIS. Thus, we next studied PKC-θ dynamics in NK cell inhibitory synapses, and how these compare to PKC-θ localization during NK cell activation. Moreover, since PKC-θ plays a role in activation of different hematopoietic cells(Baier, 1993; Hayashi, 2007) and induces SHP-1 S591 phosphorylation during T-cell activation(Liu, 2007), we tested its possible interplay with SHP-1 in NK cells. YTS-2DL1 NK cells were incubated with mCherry labeled 721.221 target cells for either 5 (early) or 20 (late) minutes (Fig. 3B). PKC-θ localization appeared dispersed at the early inhibitory NKIS and accumulated at the synapse after 20 minutes (*P*=0.0036). These dynamics were similar to SHP-1 S591 phosphorylation patterns. Furthermore, we observed PKC-θ accumulation at the activating NKIS at the initial time point, which did not significantly decrease after 20 minutes (Fig. 3 and Fig. 3-Fig. supplement 1A *P*=0.5068). This suggested that PKC-θ may play a role in regulating late inhibitory and early activating signaling pathways via SHP-1 regulation.

Next, we examined whether an interaction between PKC−θ and SHP-1 occurs in NK cells, and followed its dynamics during activation versus inhibition. To determine whether endogenous SHP-1 binds PKC-θ, YTS-2DL1 were incubated with 721.221 target cells for different time periods. Cell lysates were prepared, and immunoprecipitates (IPs) of SHP-1 were probed with anti-PKC-θ antibody. A clear elevation in the interaction between SHP-1 and PKC-θ was detected during NK cell inhibition after 20 minutes, as opposed to a relatively constant association during NK cell activation (Fig. 3-Fig. supplement 1B). Together, these data suggest that PKC-θ plays a role in human NK cell activation, and possibly in late (20 min) NK cell inhibition, potentially through SHP-1 regulation at the late inhibitory NKIS, and throughout the lifetime of the activating NKIS (5-20 minutes).

### SHP-1 phosphorylation on S591 is facilitated through PKC-θ

In order to determine whether SHP-1 S591 phosphorylation is mediated through PKC−θ in NK cells, a specific siRNA gene silencing approach was utilized in YTS-2DL1 and pNK-2DL1 cells. Cells were gene silenced for PKC−θ and incubated with 721.221 target cells (Fig. 4). Significant silencing efficiency was obtained in all experiments (*P*=0.001 and *P*=0.0135 for YTS and pNK-2DL1 cells, respectively) relative to cells transfected with non-specific (NS) siRNA control (Fig. 4A & Fig 4-Fig. supplement 1).

**Fig. 4.**
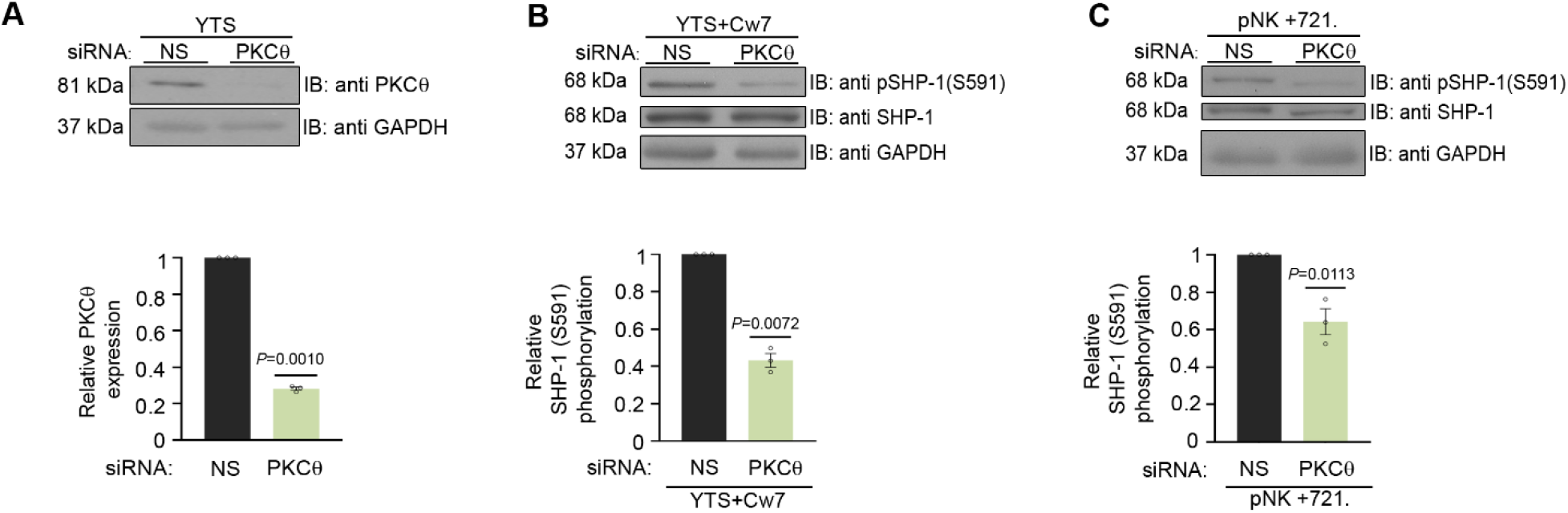
SHP-1 phosphorylation is mediated through PKC-θ. **(A)** NK cells were isolated and treated with either non-specific (control) (N.S) or PKC-θ siRNA, lysed, separated on SDS PAGE, and immunoblotted with anti-PKC-θ antibody. PKC-θ levels were measured by densitometric analysis, relative to the GAPDH loading control using ImageJ. Samples were normalized to the sample treated with N.S siRNA (*P*=0.001, quantification on the bottom showing average of 3 independent experiments). **(B)** YTS-2DL1 treated with either NS or PKC-θ siRNA were incubated with 721-HLA target cells at 37°C for 5 minutes, and cells were subsequently lysed. Lysates were separated on SDS page transferred to a nitrocellulose membrane, and immunoblotted with anti-pSHP-1 S591 antibody. SHP-1 S591 phosphorylation levels were measured by densitometric analysis, relative to the GAPDH loading control using ImageJ. Samples were normalized to the YTS-2DL1 sample treated with non-specific siRNA and incubated with 721-Cw7 target (*P*=0.0072, quantification on the bottom of independent experiments, n=3). **(C)** pNK-2DL1 cells were incubated with 721-HLA negative cells at 37°C for 5 minutes, and cells were subsequently lysed. Lysates were separated on SDS page and transferred to a nitrocellulose membrane that was immunoblotted with anti-pSHP-1 S591 antibody. SHP-1 S591 phosphorylation levels were measured by densitometric analysis, relative to the GAPDH loading control using ImageJ. Samples were normalized to the pNK-2DL1 sample treated with N.S siRNA, and incubated with 721 targets (*P*=0.0113, quantification on the bottom showing the average of 3 independent experiments). Data are shown as mean ± SEM. One sample t-tests (A, B) or one-way ANOVA with Tukey test (C) were used to calculate p values **Figure 4-source data 1 and Figure 4-Figure supplement 1-source data 1: Representative blots** **Figure 4-source data 2 Figure 4-Figure supplement 1-source data 2: Numerical data for all graphical presentation.**

Next, SHP-1 S591 phosphorylation levels were examined in YTS-2DL1 and pNK-2DL1 cells that were treated with NS siRNA vs. PKC-θ siRNA, and incubated with 721-Cw7 or 721-HLA negative target cells, in order to elucidate the role of PKC-θ on SHP-1 S591 phosphorylation during stimulation. Though S591 phosphorylation was not completely eliminated, a reduction in SHP-1 S591 phosphorylation levels could be seen in YTS-2DL1 or pNK-2DL1 treated with PKC−θ siRNA as opposed to NS siRNA (by 2.5 fold±0.03, *P*=0.0072; by 1.7 fold±0.07, *P*=0.0113, respectively) (Fig. 4B, C). Together, these data demonstrate PKC−θ is implicated in SHP-1 S591 phosphorylation in NK cells.

### PKC-θ regulates SHP-1 conformation status and its enzymatic activity at the NKIS

To determine whether SHP-1 conformation is affected by PKC-θ, YTS-2DL1 cells were co-transfected with YFP-SHP-1-CFP along with NS or PKC−θ-specific siRNA, and incubated with targets for 5 minutes. Synaptic FRET efficiency was significantly reduced in YTS-2DL1 YFP-SHP-1-CFP cells pretreated with PKC−θ siRNA versus NS siRNA (18.5±2.04% vs. 9.8±1.7%, *P*=0.0025), suggesting that the SHP-1 in the silenced cells acquires the open conformation (Fig. 5A, B). Interestingly, PKC-θ gene silencing reduced the FRET efficiency of YFP-SHP-1-CFP in the activating NKIS to similar FRET levels observed in YTS-2DL1 YFP-SHP-1-CFP cells pretreated with NS siRNA following inhibitory interactions (5.76±1.6% vs. 9.8±1.7% *P*=0.2663), (Fig. 5 A-B). Hence, although the SHP-1 conformation in activating synapses is closed and inactive, PKC−θ gene silencing increases SHP-1 open and active conformation at the activating NKIS, as detected by reduced FRET efficiency between the N’ and C’ termini of the SHP-1 sensor. This may suggest a role for PKC-θ in reducing SHP-1 activity at the activating NKIS, where SHP-1 localizes yet is inactive, and possibly during the termination of the inhibitory synapse to enable subsequent NK cell activity.

**Fig. 5.**
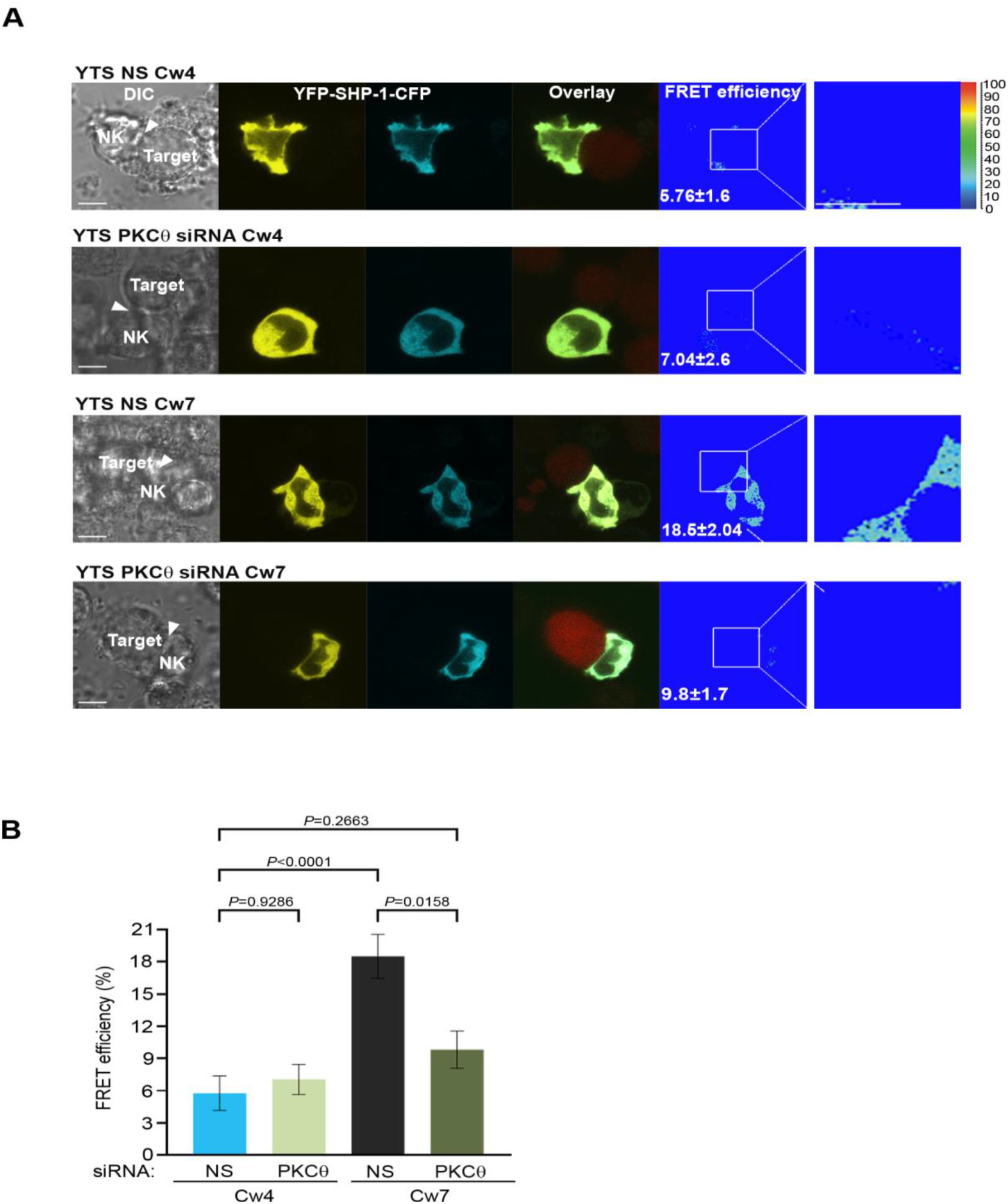
SHP-1 conformation is regulated by PKC-θ at the NKIS. **(A)** YTS-2DL1 YFP-SHP-1-CFP cells were treated with either N.S or PKC-θ siRNA and incubated over slides pre-seeded with 721-Cw4 (top panels) or Cw7 (bottom panels) target cells expressing mCherry. The cells were incubated for 5 minutes at 37°C to enable conjugate formation, and were fixed. FRET analysis was performed, as indicated. **(B)** Graph summarizing FRET efficiency during 5 min activation with 721-Cw4 or Cw7 target cells. For Cw4 NS and PKC-θ siRNA, n=52, and 63 cell conjugates were analyzed, respectively, and for Cw7 NS and PKC-θ siRNA n=67, and 61 cell conjugates were analyzed from three independent experiments, respectively. Data are shown as mean ± SEM. Two-way ANOVA with Tukey test (B) were used to calculate p values. **Figure 5-source data 1: Numerical data the graphical presentation.**

In order to examine whether the altered SHP-1 conformation induced under PKC-θ gene silencing affects SHP-1 catalytic activity, SHP-1 activity was assessed through protein tyrosine phosphatase (PTP) assay. Direct catalytic activity of SHP-1 on the pNPP substrate was shown to be higher in cells treated with PKC−θ siRNA rather than in cells treated with NS siRNA (81%±3.3% vs. 65%±3% activity, *P*=0.0246) (Fig. 5-Fig. supplement 1). Collectively, these results indicate that SHP-1 phosphorylation by PKC-θ can shift the SHP-1 conformational state from open (active) to closed (inactive), and thus reduce SHP-1 enzymatic activity.

### SHP-1 activity is modified under PKC-θ-mediated regulation

To further characterize the ability of PKC-θ to regulate SHP-1 catalytic activity, the phosphorylation profile of key signaling proteins that serve as SHP-1 substrates was compared following NK cell activation or inhibition under PKC-θ gene silencing. SHP-1 was shown to dephosphorylate VAV1 as a mechanism of terminating NK cell activation(Stebbins, 2003). Furthermore, our recent studies identified novel SHP-1 substrates including PLCγ1/2 and LAT(Matalon, 2016).

We therefore examined how PKC-θ gene silencing affects SHP-1 catalytic activity on its substrates in YTS and pNK-2DL1 cells. In order to assess the phosphorylation status of these proteins, cells were treated either PKC−θ siRNA or with NS siRNA and incubated with inhibitory Cw4-721 or activating Cw7-721 or 721-HLA negative target cells. IB of pVAV1 (Y160) revealed an approximately 2-fold reduction in phosphorylation levels in activated YTS-2DL1 samples treated with PKC-θ siRNA in contrast to NS siRNA (*P*=0.0001) (Fig. 6A). Similarly, pNK-2DL1 cells demonstrated a similar reduction in pVAV1 (Y160) phosphorylation levels subsequent to PKC-θ gene silencing (*P*=0.0192) (Fig. 6A). Furthermore, phosphorylation of VAV1 in activating interactions following PKC−θ gene silencing was similar to its phosphorylation during inhibitory interactions (YTS and pNK-2DL1 cells treated with NS siRNA). In addition to VAV1, pNK-2DL1 cells showed a 1.8 fold±0.2 reduction in pPLCγ1 levels (*P*=0.0032) when treated with PKC−θ siRNA (Fig. 6B). A similar trend can be seen in YTS-2DL1 cells in which IP of PLCγ1 and IB of pPLCγ1 (Y783) revealed a reduction of 1.4 fold±0.04 in pPLCγ1 levels following PKC-θ gene silencing under activating interactions (P=0.0074) (Fig. 6-Fig. supplement 1). Together these data suggest that PKC-θ-mediated regulation of SHP-1 catalytic activity may promote SHP-1 inactivation, and thus enhance NK cell reactivity.

**Fig 6.**
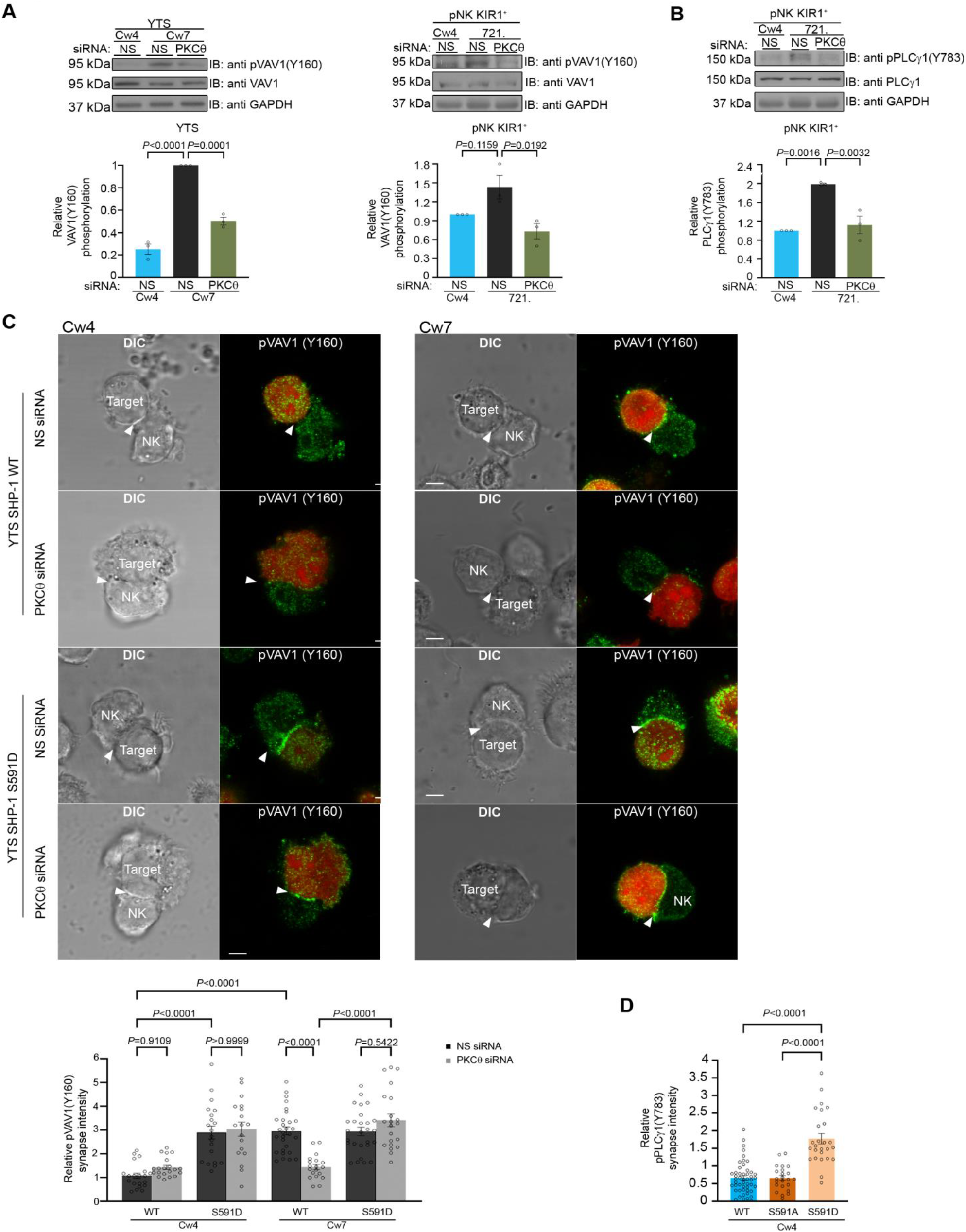
SHP-1 substrate phosphorylation is reduced following PKC-θ silencing. **(A)** YTS-2DL1 or pNK-2DL1 cells were transfected with 500 pmol of PKC-θ siRNA for 48hrs. Cells were incubated with target cells for 5 minutes at 37°C and then lysed. Lysates were separated on SDS page and immunoblotted with anti pVAV-1 (Y160) antibody. Phosphorylation levels were measured by densitometric analysis, relative to the GAPDH loading control using ImageJ. Samples were normalized according to the YTS-2DL1 or pNK-2DL1 NS siRNA Cw4 sample. Quantification of independent experiments is shown on the bottom; n=3 for YTS and n=4 for pNK experiments. **(B)** pNK-2DL1 cells were transfected with either N.S or PKC−θ siRNA 48 hrs prior to each experiment. Cells were incubated with either 721-Cw4 or 721-HLA negative target cells for 5 minutes at 37°C, and subsequently lysed. Lysates were separated on SDS page and immunoblotted with anti pPLCγ-1 (Y783) antibody. Phosphorylation levels were measured by densitometric analysis, relative to the GAPDH loading control using ImageJ. Samples were normalized to the pNK-2DL1 NS siRNA Cw4 sample. Quantification of independent experiments is shown on the bottom; n=3. The blot in (B) was stripped and reblotted against pVAV-1(Y160) in (A). **(C)** YTS-2DL1 WT or YTS-2DL1 S591D cells were transfected with either NS or PKC-θ siRNA and incubated over slides pre-seeded with 721-Cw4 or Cw7 target cells expressing mCherry. The cells were incubated for 5 minutes at 37°C to enable conjugate formation, and fixed. pVAV-1 (Y160) was subsequently labeled with primary rabbit anti-pVAV-1 (Y160) antibody and secondary anti-rabbit 488 antibody. Quantification is shown on the bottom; for YTS-2DL1 WT NS or PKC-θ siRNA vs. Cw4 and Cw7, n=21 and 28 cell conjugates were analyzed, respectively. For YTS-2DL1 S591D NS or PKC-θ siRNA vs. Cw4 and Cw7, n=21 and 28 cell conjugates were analyzed from three independent experiments, respectively. **(D)** YTS-2DL1 SHP-1^-/-^ cells were transfected with either WT YFP-SHP-1, S591A YFP-SHP-1, or S591D YFP-SHP-1 phosphorylation mutants and incubated on slides with mCherry expressing 721-Cw4 target cells at 37°C. After 5 min incubation, the cells were fixed and stained with anti-pPLCγ-1 (Y783) and secondary Alexa-488 antibody; accumulation of phosphorylated PLCγ-1 at the NKIS was determined. NK cells were distinguished from targets based on mCherry expression by the target cells. Graph summarizes the relative synapse staining intensities. For WT YFP-SHP-1, S591A YFP-SHP-1, and S591D YFP-SHP-1, n= 48, 48, and 25 cell conjugates from three independent experiments analyzed, respectively. Data are shown as mean ± SEM. One-way ANOVA with Tukey test (A, B, D) or two-way ANOVA with Tukey test (C) were used to calculate p values **Figure 6-source data 1 and Figure 6-Figure supplement 1-source data 1: Representative blots** **Figure 6-source data 2 and 3 Figure 6-Figure supplement 1-source data 1: Numerical data for all graphical presentation.**

In order to demonstrate that PKC-θ regulation of SHP-1 via S591 influences SHP-1 substrate phosphorylation and activation, and not PKC-θ silencing per-se, a mutant YTS-2DL1 line expressing a SHP-1 mutant that mimics the constitutively phosphorylated state, SHP-1 S591D, was created utilizing CRISPR/CAS9, as previously reported (Ben-Shmuel, 2020). In cells expressing this mutant, PKC-θ silencing would not be expected to impact SHP-1 activity(Egelhoff *et al*., 1993; Huang *et al*., 1994; Léger *et al*., 1997). Hence, we expected that the S591D mutant would demonstrate high accumulation of pVAV1 in the synapse, irrespective of PKC-θ silencing. WT YTS-2DL1 or YTS-2DL1 SHP-1 S591D NK cells were treated with either NS or PKC-θ siRNA, and incubated with mCherry-expressing target cells to assess pVAV1 Y160 synaptic accumulation. As expected, pVAV1 Y160 accumulation was observed during activating but not inhibitory NK cell interactions (*P*<0.0001), and in accordance with our results, PKC-θ siRNA treated NK cells had a 2±0.2 fold reduction in pVAV1 recruitment to the activating NKIS (*P*<0.0001), indicating increased SHP-1 activity in the absence of PKC-θ under activating interactions (Fig. 6C). No significant decrease was observed, however, in pVAV1 recruitment in SHP-1 S591D NK cells treated with either NS or PKC-θ siRNA under either activating or inhibitory interactions (*P*=0.9109, *P*=0.9999, respectively). Furthermore, YTS-2DL1 SHP1^-/-^ cells were reconstituted with either YFP-SHP-1 WT or with YFP-SHP-1 mutant constructs, including the constitutively phosphorylated SHP-1 mimetic (YFP-SHP-1 S591D, constitutively inactive) or an SHP-1 phospho-abolishing mutant (YFP-SHP-1 S591A, constitutively active). YTS-2DL1 SHP-1^-/-^ cells transfected with YFP-SHP-1 were incubated with Cw4-721 targets expressing mCherry, and stained for pPLCγ1 (Y783) (Fig. 6D). We previously showed that pPLCγ1 (Y783) accumulation at the NKIS is favored during activating rather than inhibitory NK cell interactions^14^. Interestingly, cells expressing SHP-1 S591D showed high levels of pPLCγ1 (Y783) accumulation at the NKIS, even though the NKIS was inhibitory. These levels were approximately 3 fold higher compared to YTS-2DL1 cells expressing SHP-1 WT or SHP-1 S591A (*P*<0.0001). Collectively, these results demonstrate that PKC-θ negatively regulates SHP-1 activity by phosphorylating S591, as demonstrated through higher VAV1 and PLCγ1 tyrosine phosphorylation, and accumulation at the NK synapse.

### PKC-θ-mediated regulation of SHP-1 augments NK cell activation

SHP-1 tunes NK cell activation, and, as we demonstrated here, PKC-θ-mediated phosphorylation of SHP-1 impacts its enzymatic activity. We therefore next determined the effect of PKC-θ regulation on NK cell effector function and activation. PKC-θ silencing in NK cells resulted in a decrease in the phosphorylation profile of VAV1 and PLCγ1 in a SHP-1 dependent manner (Fig. 6). These data suggest that the PKC−θ:SHP-1 axis may affect NK cell activation and cytotoxicity, in a manner that is highly dependent on calcium flux and actin reorganization(Orange *et al*., 2002; Stebbins, 2003; Bryceson *et al*., 2006; Caraux *et al*., 2006; Upshaw *et al*., 2006; Andzelm *et al*., 2007; Kim *et al*., 2010; Dong *et al*., 2012; Mizesko *et al*., 2013; Carisey *et al*., 2018). To this end, PKC−θ was gene-silenced in YTS-2DL1 cells, and intracellular calcium flux was measured in NK cells incubated with activating 721-Cw7 target cells. As expected, calcium flux levels were higher in activated vs. inhibited NK cells treated with NS siRNA (black vs. blue curve, Fig. 7A). However, PKC-θ siRNA reversed this trend, reducing calcium flux levels during NK cell activation near the levels obtained during NK cell inhibition (green vs. blue curve, Fig. 7A). Similar results were obtained in pNK-2DL1cells transfected with NS or PKC-θ siRNA and incubated with activating 721-HLA negative or 721-Cw4 expressing cells (Fig. 7-Fig. supplement 1). These results are consistent with the reduced phosphorylation levels of the calcium regulator, PLCγ1 (Fig. 6B and D and Fig. 6-Fig. supplement 1).

**Fig 7.**
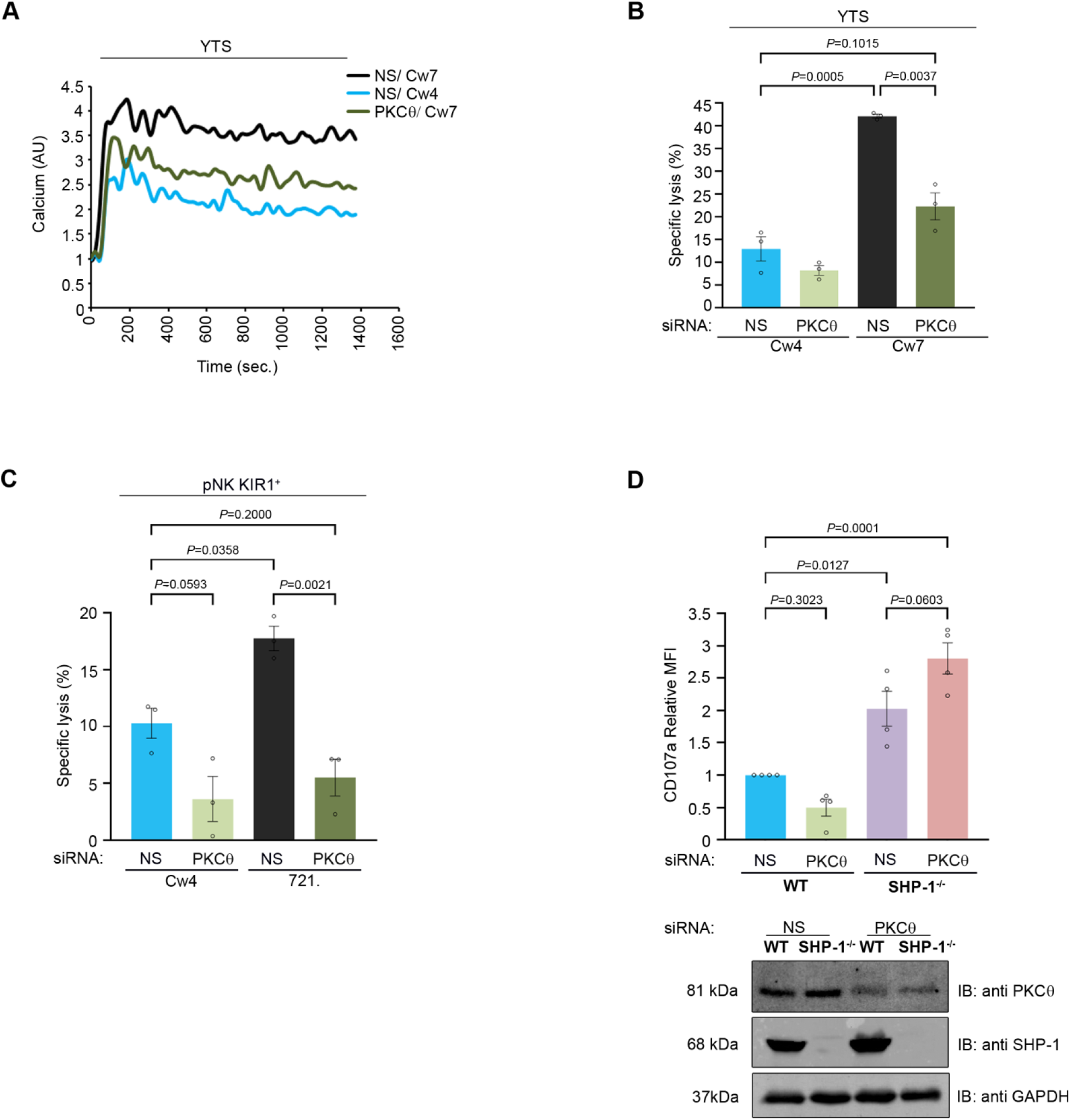
NK cell activation threshold is increased following gene silencing of PKC-θ. YTS-2DL1 cells were transfected with specific PKC-θ siRNA or with N.S siRNA and used after 48 hrs. **(A)** YTS-2DL1 cells were loaded with calcium-sensitive Fluo-3-AM, and analyzed for basal intracellular calcium levels for 1 minute. The NK cells were then mixed with 721-Cw4 or Cw7 target cells and incubated at 37°C and analyzed by spectrofluorometry. **(B)** YTS-2DL1 cells were incubated with [^35^S]Met labeled 721-Cw4 or Cw7 target cells at a ratio of 10:1 for 5 hours at 37°C. The specific lysis of target cells was measured. The graph summarizes 3 independent experiments. **(C)** pNK-2DL1 cells were incubated with [^35^S]Met labeled 721-Cw4 or HLA negative target cells at a ratio of 10:1 for 5 hours at 37°C. The specific target cell lysis was measured. The graph shows the average of 3 independent experiments. **(D)** WT YTS-2DL1 or YTS-2DL1 SHP-1^-/-^ cells were treated with either NS or PKC−θ specific siRNA 24 hours before incubation with mCherry-expressing 721-Cw4 target cells for 2 hours at 37°C, and analyzed for degranulation via FACS relative to the WT NS treated cells incubated with 721-Cw4 targets, as described in the Materials and Methods. A representative blot is shown (bottom) demonstrating PKC-θ and SHP-1 expression; the graph shows the average of 4 independent experiments. Data are shown as mean ± SEM. Two-way ANOVA with Tukey test (B, C) or one-way ANOVA with Tukey test (D) were used to calculate p values. **Figure 7-source data 1 and Figure 7-Figure supplement 1-source data: Numerical data for all graphical presentation.** **Figure 7-source data 2: Representative blots**

To assess how the PKC-θ:SHP-1 regulation controls NK cell cytotoxic potential, YTS-2DL1 cells were gene silenced for PKC-θ and incubated with 721-Cw4 or Cw7 target cells, and target cell lysis was measured through Met release assay [^35^S] a direct measurement of NK cell killing of cancerous cells (Fig. 7B). Target cell death was significantly reduced in YTS-2DL1 samples that were gene silenced for PKC-θ under activating interactions (22%±3% vs. 42%±0.4%, *P*=0.0037). In fact, activating target cells (Cw7) incubated with PKC-θ-silenced NK cells demonstrated lysis levels similar to those of inhibitory targets (Cw4) incubated with YTS-2DL1 cells treated with NS siRNA (22%±3% vs. 13%±2.7%, *P*=0.1015). These data were verified in human primary NK cells treated with NS or PKC-θ siRNA (5.5%±1.6% vs. 18%±1.06, *P*=0.0021, and 5.5%±1.6% vs. 10%±1.32%, *P*=0.2) (Fig. 7C and Fig. 7-Fig. supplement 1). To verify that absence of PKC-θ influences NK cell effector functions through SHP-1 activity, and that the effects of PKC-θ silencing are abrogated in the absence of SHP-1, SHP-1^-/-^ NK cells were additionally gene silenced for PKC−θ (Fig. 7D). Subsequently, WT and SHP-1^-/-^ NK cells either treated with NS or PKC−θ siRNA were subjected to CD107a degranulation assay. As can be seen in Fig. 7D, NK cells that were deficient in SHP-1 or both SHP-1 and PKC-θ displayed higher degranulation than WT cells (*P*=0.0127 and *P*=0.0001), and were not significantly influenced by the presence or absence of PKC−θ (*P*=0.0603). Taken together, these data demonstrate that PKC-θ negatively regulates SHP-1 conformational state and activity, thereby serving as a mechanism for maintaining NK cell activation and cytotoxic potential.

### SHP-1 deficient NK cells are unaffected by PKC-θ silencing, and promote superior anti-tumor clearance relative to WT cells

We recently demonstrated that abrogation of SHP-1 activity in engineered NK cells can have enhanced benefits for NK-based immunotherapeutic approaches(Ben-Shmuel, 2020). To verify that NK cells silenced for PKC−θ have lower anti-tumor cytotoxicity due to enhanced SHP-1 activity, and not due to absence of PKC-θ per se, NOD-Rag1^null^IL2rg^null^ (RAG) immunodeficient mice were subcutaneously engrafted with 721-Cw4 /Cw7 cells and injected with either 3×10^6^ WT YTS-2DL1, WT YTS-2DL1 that were treated with PKC-θ siRNA, YTS-2DL1 SHP-1 deficient cells(Matalon, 2018) or YTS-2DL1 SHP-1 deficient cells that were treated with PKC-θ siRNA, every 3 days for a total of six treatments. Tumor volumes were monitored daily to examine the effect on tumor size and average growth rate. If PKC-θ affects NK cell cytotoxicity in a dominant alternative pathway to SHP-1, then the observed phenotype of SHP-1 silenced/mutated NK cells (highly increased killing and perturbation of tumor growth) would be expected be abrogated, at least moderately, upon PKC-θ silencing. Indeed, mice injected with YTS-2DL1 SHP-1 deficient/catalytically inactive cells that were treated with PKC−θ siRNA demonstrated slower tumor growth rate and smaller tumor volumes compared to mice injected with WT NK cells (Fig. 8). Thus, regulation of SHP-1 through PKC-θ impacts NK cell cytotoxicity and affects NK cell capability for in-vivo tumor clearance.

**Fig. 8.**
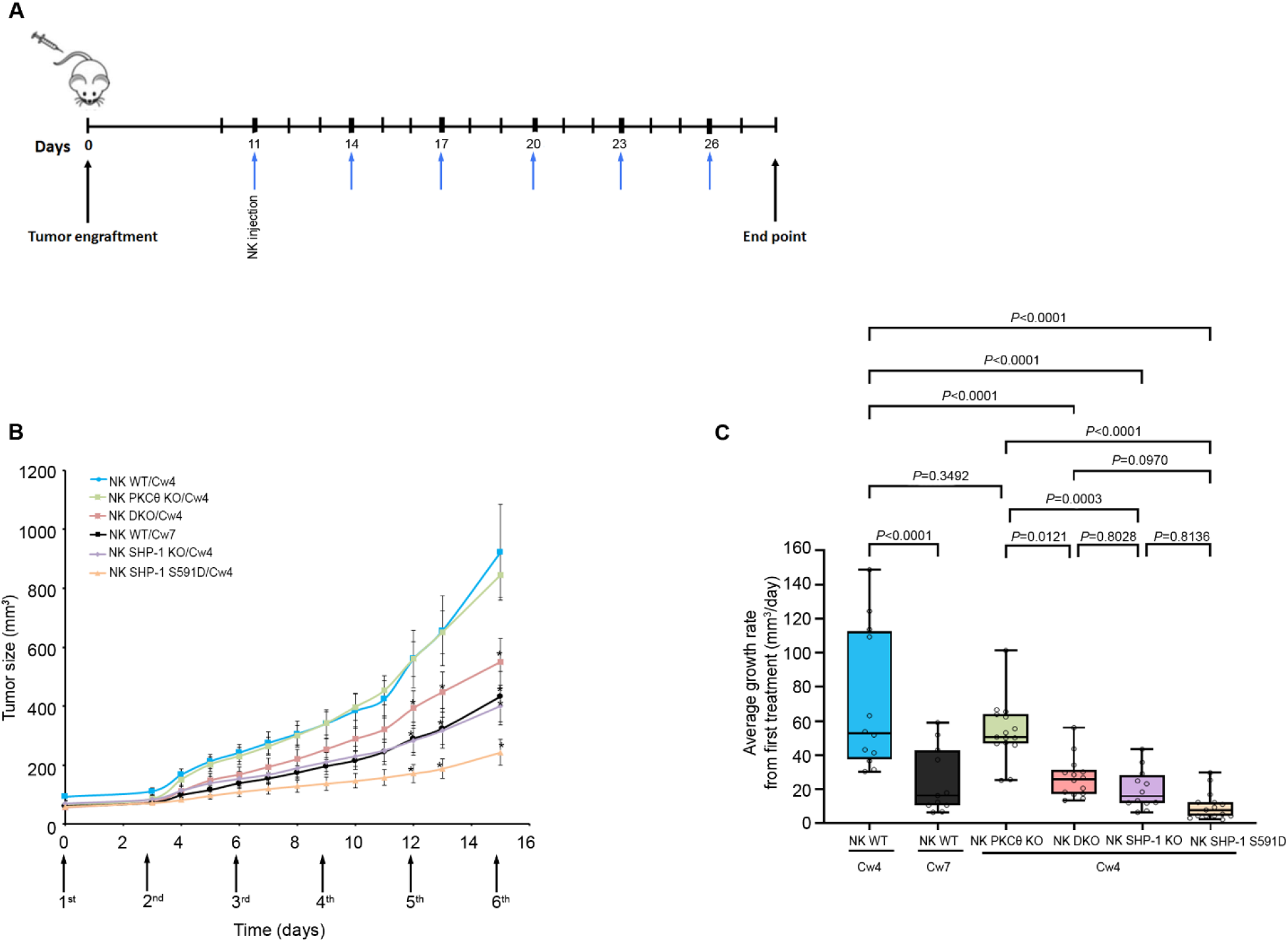
SHP-1 deficient NK cells exhibit enhanced anti-tumor activity in vivo, irrespective of PKC-θ expression. (A) Schematic representation of the experimental timeline. **(B)** Tumor volumes measured daily in NOD-Rag1^null^IL2rg^null^ (RAG) mice. NOD-Rag1^null^IL2rg^null^ (RAG) mice were subcutaneously injected with 3×10^6^ 721-Cw4 or Cw7 expressing tumor cells. Mice were administered by intratumor injections every 3 days with either 3*10^6^ irradiated WT YTS-2DL1, WT YTS-2DL1 that were treated with PKC-θ siRNA (NK PKC-θ KO), YTS-2DL1 SHP-1 deficient cells (NK SHP-1 KO) or YTS-2DL1 SHP-1 deficient cells that were treated with PKC-θ siRNA (NK DKO). (C) Average growth rates of tumors from the first treatment to the end point. Tumor volumes and tumor growth rates were calculated as described in the Methods. Data are shown as mean ± SEM. Two-way ANOVA with Tukey test (B, C) were used to calculate p values. **Figure 8-source data 1: Numerical data for all graphical presentation.**

## Discussion

SHP-1 plays a central role in the regulation of NK cell activation threshold(Stebbins, 2003; Lanier, 2005; Long, 2008; Matalon, 2016) and is critical for the NK cell education process(Viant *et al*., 2014). Importantly, activation of SHP-1 and its homolog SHP-2 (an additional critical inhibitory NK cell regulatory enzyme(Purdy *et al*., 2009)) are the major mechanisms operating downstream to immune checkpoint receptors, such as PD-1, CTLA-4, BTLA, and LAG-3(Lee *et al*., 1998; Watanabe *et al*., 2003; Sheppard *et al*., 2004; Huang *et al*., 2015). It is therefore of great interest to understand how this important regulator is kept in check during NK cell activation and inhibition, as this regulatory mechanism may have critical consequences for additional immune cells, immune checkpoint pathways, and immunotherapies.

NK cell activity is regulated by a balance between stimulatory or inhibitory signals, yet how these signals are integrated, and how intracellular NK cell activation is maintained and controlled through these multiple receptors remains incompletely understood. Moreover, the presence of SHP-1 at the activating NKIS and rapid dispersal from the inhibitory NKIS(Vyas, 2002a) suggest as yet unknown roles and regulatory mechanisms. Here, we elucidate these unresolved mechanisms by showing that the NK cell activation threshold is maintained through downregulation of SHP-1 phosphatase activity regulated by PKC−θ.

Studies published on outcomes of SHP-1 C’ terminal serine phosphorylation in non-NK cell systems demonstrated contradicting results(Li, 1995; Brumell *et al*., 1997; Jones, 2004; Poole, 2005; Liu, 2007). Here, we demonstrate that phosphorylation of SHP-1 by PKC-θ during NK cell activation on serine 591 maintains heightened NK cell effector function by retaining the SHP-1 closed and inactive conformation. This mechanism preserves SHP-1 substrate phosphorylation, and consequently sustains calcium flux and cytotoxicity. Thus the PKC-θ: SHP-1 axis may play an important role in NK immune surveillance and tumor clearance, a topic that has recently gained much attention in the immunotherapy field, especially in the context of adoptive cell transfer and genetically engineered NK cells(Dotta *et al*., 2007; Morandi *et al*., 2008; Ahern *et al*., 2011; Jochems *et al*., 2016; Siegler *et al*., 2017).

Our findings may answer an important question regarding how SHP-1 activity is down modulated after dissociation of the inhibitory NKIS, and how its signaling is downregulated during cytolytic NK responses. Phosphorylation of SHP-1 during termination of the inhibitory NKIS may retain the activity of NK cells to subsequent targets, whereas its immediate phosphorylation at the activating NKIS maintains the NK cell lytic capacity. This is consistent with the SHP-1 conformation patterns shown here (Fig. 2) within the activating and inhibitory NKIS.

Interestingly, mathematical modeling analysis in NK cells suggest that weak ligands for activating receptors (bearing partially phosphorylated ITAMs) can recruit SHP-1 and thereby increase the NK activation threshold, while under certain conditions (such as low concentration of SHP-1 and weak activating ligands) inhibitory receptors can aid in NK cell activation(Das, 2010). Thus, it is possible that one mechanism facilitating target cell escape from NK cell lysis is selection for cells expressing weak activating ligands which induce SHP-1 recruitment. It is clear that the transition of SHP-1 between inhibiting and activating receptors must be tightly regulated to ensure proper NK cell responses.

PKC−θ is the only PKC isoform that was shown to accumulate at T-cell and NK cell synapses(Merino, 2012),(Monks *et al*., 1997). In T cells, it is clear that PKC-θ plays important role in T cell activation and survival by activating several downstream pathways, including NFkB and AP-1 as major targets(Wang *et al*., 2015). In addition, Merino et al previously showed that clustering of PKC-θ at the NKIS amplifies murine NK cell activation and effector functions(Merino, 2012), but the mechanism by which it exerts this function and specifically in human cells remained mostly unknown. Several studies suggest the involvement of PKC-θ in signal transduction, anti-tumoral activity and NK cell degranulation(Anel, Juan I. Aguiló, *et al*., 2012). Here we demonstrate that in human NK cells, PKC-θ phosphorylates SHP-1 on S591 (Fig. 4, Fig. 3-Fig. supplement 1C), suppresses SHP-1 activity, and thereby increases NK cell activity. PKC-θ appears to regulate SHP-1 activity during the late inhibitory synapse and through the full duration of activating NK: target cell interactions. Indeed, gene silencing of PKC-θ resulted in the closed SHP-1 conformation (Fig. 5), implying that SHP-1 favors an open and active conformation in the absence of PKC−θ. This conformational change in SHP-1 also impacts its catalytic activity. These findings were strengthened through *in-vivo* experiments, which demonstrated that absence of SHP-1 increased NK cell activity and decreased tumor progression, irrespective of PKC-θ expression, implying that PKC-θ indeed maintains NK cell activation through its regulation of SHP-1.

Several studies show that deficiency or inhibition of PKC-θ could potentially decrease the severity of autoimmunity, allergy and chronic inflammation. For example in animal models of intestinal inflammatory disease (chronic colitis), PKC-θ KO mice showed decreased T cell proliferation and cytokines production(Chand *et al*., 2012; Curnock *et al*., 2014; Nicolle *et al*., 2021) suggesting that PKC-θ inhibitors could be useful as a therapeutic approach for inflammatory disorders(Chand, 2012; Curnock, 2014).

Though pSHP-1 levels were substantially reduced after PKC-θ gene silencing, basal levels were still evident. It is possible that other serine/threonine kinases or isoforms such as PKC−α may also be involved in this process(ϑονεσ, 2004). Another possible mechanism may involve PKC−θ cross talk with the NK cytoskeleton. We recently demonstrated that SHP-1 conformation and activity is dependent on association to the cytoskeletal machinery and cytoskeletal dynamics(Matalon, 2018). One possibility is that crosstalk between PKC−θ and SHP-1 mediates actin: SHP-1 binding or disassociation, thereby regulating SHP-1 catalytic activity and conformation. Furthermore, PKC−θ phosphorylates the Wiskott-Interacting Protein (WIP) in NK cells upon activation, and this facilitates recruitment of actin and myosin IIA to WIP and WASp for formation of the multi protein complex(Krzewski *et al*., 2006). Hence PKC-θ activity could affect acto-myosin dynamics and velocity at the NKIS, influencing SHP-1 activity. It would thus be interesting to elucidate how the interaction of SHP-1 with the cytoskeletal machinery changes as a function of PKC-θ activity.

After SHP-1 mediates its function and prevents NK cell activity during an inhibitory interaction, its activity should be downregulated to enable subsequent NK cell interactions with activating susceptible targets. This can be seen by late phosphorylation (after 20 minutes) of SHP-1 S591 during inhibitory NK cell synapses (Fig. 1), and may serve as a mechanism of kinetic priming, ensuring that NK cell activity remains stable for future contacts(Choi, 2013).

Recently it was shown that as NK cells are constantly situated in an environment that requires simultaneous killing of target cells and sparing of others, accurate and specific convergence of lytic granules aids in killing only relevant targets, while sparing healthy cells(Hsu, 2016). Moreover, a study by Netter et al demonstrated how initiation and termination of the activating NKIS is highly regulated, and enables serial NK cell killing by accelerating detachment from one target cell and simultaneous formation of new cytotoxic NK cell synapses(Netter, 2017). Additional reports by Srpan et al. describe NK cell serial killing via shedding of surface receptors(Srpan, 2018). Thus, sequential and highly regulated signaling is needed for NK cell maneuverability in a complex immune environment. PKC-θ;SHP1 regulation may enable an NK cell to rapidly survey its surroundings and maintain its effector functions in a complex immune environment that requires simultaneous sparing of normal cells, and killing of malignant targets. This mechanism would also ensure that NK cell effector functions are maintained in the presence of an activating NK cell synapse. Cancer cells can express MHC-I and other inhibitory NK cell ligands; thus, recruitment and activation of PKC-θ at the activating NKIS may limit SHP-1 association to ITIMs that could prevent tumor cell killing. It would be interesting to assess the effect of S591 phosphorylation on SHP-1:KIR ITIM association, as this phosphorylation may not only affect the NK cell activation threshold, but also NK cell education, as SHP-1 was shown to be vital for regulating NK cell responsiveness to tumors(Viant, 2014).

Due to the fact that NK cells are also implicated in auto immune disorders such as multiple sclerosis(Morandi, 2008), rheumatoid arthritis(Ahern, 2011), and type I diabetes (Dotta, 2007), deregulation of SHP-1 activity may be a factor affecting normal NK cell functions against healthy cells. SHP-1 was also shown to serve as an important transcription factor that impacts cell proliferation and survival. Some studies examined its role in tumor transformation(Wu *et al*., 2003). It would be interesting to examine whether mutations or regulation of SHP-1 S591 or other known or as yet unidentified residues may impact its function. Understanding regulatory protein circuits in NK cells will be an important step for further elucidating complex pathways such as NK signaling cascades and NK cell education, and potentially open doors to innovative immunotherapies shaping NK cell behavior.

## Materials and Methods

### Cell lines and reagents

The following cells were used in this study: YTS NK cell line expressing the inhibitory KIR2DL1 (referred as YTS-2DL1), B-cell lymphoma, 721.221 cells (referred as 721) expressing no HLA, and 721 cells expressing either HLA-Cw4 or –Cw7, and K562. These cells were kindly provided by Prof. Ofer Mandelboim (Department of Microbiology and Immunology, Faculty of Medicine, Hebrew University of Jerusalem, Israel). YTS cells were cultured in Iscove’s medium supplemented with 10% fetal bovine serum (FBS), 2mM L-glutamine, 50μg/ml penicillin, 50μg/ml streptomycin and 50μM 2-mercapto-ethanol. 721.221 and K562 cells were cultured in RPMI supplemented with 10% FBS, 2mM L-glutamine, 50μg/ml penicillin, 50μg/ml streptomycin, 1% non-essential amino acids, and 1% sodium pyruvate.

### Antibodies

Antibodies and their sources were as follows: Mouse anti-PLCγ1 (Upstate), Mouse anti-VAV1 (D7), Rabbit anti-SHP-1 and Mouse anti-GAPDH (Santa Cruz), Rabbit anti-pSHP-1 (S591) (ECM Biosciences), Rabbit anti-pPLCγ1 (Y783) (Bio Source), Rabbit anti-pVAV1 (Y160) (Bio Source), and Mouse anti-PKC-θ (Santa Cruz). Secondary antibodies: Goat anti-Mouse (Sigma-Aldrich), Goat anti-Rabbit (Santa Cruz).

### Primary NK cells

Primary NK cells were isolated from peripheral blood lymphocytes (PBLs) of healthy donors using the EasySep™ human NK Cell enrichment kit (STEMCELL Technologies). Subsequently, KIR2DL1 expressing cells were isolated by staining the entire NK cell population with anti-KIR1-PE antibody (Miltenyi Biotec) followed by magnetic separation using the EasySep™ human PE selection kit (STEMCELL Technologies) according to the manufacturer’s instructions. NK cell isolation efficiencies were >95%. The NK cells were plated in 96-well U-bottomed plates and grown in the presence of irradiated PBLs from two donors (5×10^4^ cells from each donor per well) as feeder cells. Cells were expanded in a complete medium containing 1µg/ml of PHA, and 400 U/ml rhuIL-2 (Prospec). Before experiments, cells were washed to remove the PHA and IL-2, and cultured 48 hours in 60% Dulbecco’s modified Eagle medium (DMEM) and 25% F-12 medium supplemented with 10% human serum, 2mM L-glutamine, 50μg/ml penicillin, 50μg/ml streptomycin, 1% non-essential amino acids and 1% sodium pyruvate.

### CRISPR/CAS9 mediated gene knockdown and S591D point mutation

CRISPR/CAS9 knockdown of SHP-1 in YTS cells was conducted according to published protocol and was performed as described in detail (Ran *et al*., 2013; Matalon, 2018; Ben-Shmuel, 2020). The pSpCas9(BB)-2A-GFP (PX458) vector was purchased from Addgene (plasmid # 48138). RNA guide sequences targeted to the SHP-1 locus and for knock-in of the S591D mutation were constructed using an online CRISPR design tool (Zhang Lab).

### RNA interference

Small interfering RNA (siRNA) to human 3’ UTR PKC-θ was purchased from Sigma Aldrich. YTS-2DL1 cells were transfected with siRNA specific for the indicated genes, or non-specific (NS) siRNA as a control, using an AMAXA electroporator.

For knockdown of PKC-θ gene expression, pools of independent specific siRNA oligonucleotides were as follows:

5’ CUCUUCACCUGGGCGCCAADTDT 3’

5’ UUGGCGCCCAGGUGAAGAG 3’

Non-specific siRNA: Pools of nontargeting (nonspecific), negative control siRNA duplexes were obtained from Dharmacon with the following sequences: 5’ UAGCGACUAAACACAUCAA 3’,

5’UAAGGCUAUGAAGAGAUAC3’, 5’AUGUAUUGGCCUGUAUUAG3’,5’ AUGAACGUGAAUUGCUCAA 3’, and 5’UGGUUUACAUGUCGACUAA3’

### Cellular imaging by confocal microscopy

Dynamic fluorescence and differential interference contrast microscopy (DIC) images of NK-target conjugates were collected using a Zeiss 510 Meta confocal microscope. All images were collected using a 63 X Plan-Apochromat objective (Carl Zeiss).

Chambered cover slips (LabTek) were cleaned by treatment with 1 M HCl, 70% ethanol for 30 min and dried at 60^0^C for 30 min. The chambers were treated with a 0.01% (wt/vol) poly-L-lysine solution (Sigma) for 5min, drained, and dried at 60^0^C for 30 min.

For NK-target conjugation assays, 5×10^5^ target cells were seeded over the bottom of the chamber in 300μl Optimem medium for 2 hours at 37°C, after which nonadherent cells were rinsed. Then, 5×10^5^ NK cells were seeded over the chambers, containing imaging buffer (RPMI medium with 25mM HEPES without phenol red or serum), and allowed to form conjugates with the target cells for the indicated times at 37°C. Following activation, the cells were fixed for 30 min with 2.5% paraformaldehyde, and washed twice with PBS. The NK and target cells in the conjugates were distinguished according to fluorescence signal, with the target cells expressing mCherry. For evaluation of phosphoprotein accumulation at the NKIS, cells were permeabilized with 0.1% Triton X-100 for 5 min. Cells were then blocked for 1 hour in PFN buffer (PBS without Ca2+ and Mg2^+^ and containing 10% FBS and 0.02% azide) with 2% normal goat serum (Jackson ImmunoResearch). Cells were incubated for 1 hour with the appropriate primary antibodies diluted in blocking medium, followed by staining with isotype-specific, Alexa Fluor–conjugated antibodies for 30 min. Cells were washed three times with PFN between steps. The relative fluorescence intensities of the proteins at the NKIS were determined by measuring the ratio between the fluorescence intensity at the NKIS, relative to that at a non-NKIS site using ImageJ software.

### Image processing and quantification

The acquired images were extracted with the LSM browser (Carl Zeiss), cropped and composed into Figures using Adobe Photoshop.

### FRET analysis

FRET was measured by the donor-sensitized acceptor fluorescence technique as previously described(Barda-Saad, 2005; Pauker, 2012; Fried, 2014). Briefly, three sets of filters were used: one optimized for donor fluorescence (excitation, 458 nm, and emission, 465 to 510 nm), a second for acceptor fluorescence (excitation, 514 nm, and emission, 530 to 600 nm), and a third for FRET (excitation, 458 nm; emission, 530 to 600 nm).

### FRET correction

FRET correction was performed as described in detail(Barda-Saad, 2005; Pauker, 2012; Fried, 2014). The non-FRET components were calculated and removed using calibration curves derived from images of single-labeled CFP- or YFP-expressing cells. Sets of reference images were obtained using the same acquisition parameters as those used for the experimental FRET images. To correct for CFP “bleed through,” the intensity of each pixel in the CFP image from CFP-expressing cells was compared to the equivalent pixel in the FRET image of the same cells. A calibration curve was derived to define the level of CFP fluorescence seen in the FRET image as a function of the fluorescence in the CFP image. A similar calibration curve was obtained defining the amount of YFP fluorescence appearing in the FRET image as a function of the intensity in the YFP image, using images of cells expressing only YFP. Separate calibration curves were derived for each set of acquisition parameters used in the FRET experiments. Then, using the appropriate calibration curves, together with the CFP and YFP images, the amount of CFP bleed through and YFP cross excitation was calculated for each pixel in the experimental FRET images. These non-FRET components were subtracted from the raw FRET images, yielding corrected FRET images.

### FRET efficiency calculation

The FRET efficiency (FRETeff) was calculated on a pixel-by-pixel basis using the following equation: FRETeff = FRETcorr/(FRETcorr + CFP) X 100%, where FRETcorr is the pixel intensity in the corrected FRET image, and CFP is the intensity of the corresponding pixel in the CFP channel image.

To increase the reliability of the calculations and to prevent low-level noise from distorting the calculated ratio, we excluded pixels below 50 intensity units and saturated pixels from the calculations and set their intensities to zero. These pixels are shown in black in the “pseudocolored” FRET efficiency images.

To estimate the significance of the FRET efficiency values obtained, and to exclude the possibility of false-positive FRET results, we prepared cells expressing free CFP and free YFP as negative controls. The FRET efficiency in the negative-control system was measured and calculated in the same way as in the main experiment. FRET efficiency values obtained from the negative-control samples were subtracted from the values obtained in the main experiments. Image processing and measurements were performed using IPLab software, version 3.9.

### PTP Assay

SHP-1 catalytic activity was determined by measuring the hydrolysis of the exogenous substrate p-Nitrophenyl Phosphate (pNPP) by SHP-1, as previously described(Lorenz, 2011; Matalon, 2018). NK cells (2-5 x 10^6^) were incubated with target cells at ratio of 1:1 at 37°C for 5 min before lysis. Cells were lysed with ice-cold passive lysis buffer (1.25% Brij, 0.625% n-Octyl-b-D-glucoside, 31.3 mM Tris–HCl, pH 7.4, 150 mM NaCl, 6.25mM ethylenediaminetetraacetic acid, and complete protease inhibitor tablets (Roche)). Cell lysates were subjected to IP with anti–SHP-1 antibody. Immunoprecipitates were washed twice with ice-cold passive washing buffer (0.1% Brij, 50 mM Tris–HCl, pH 7.4, 300mM NaCl, and 3.75mM ethylenediaminetetraacetic acid), and three times with phosphatase buffer (150mM NaCl, 50mM HEPES, 10mM ethylenediaminetetraacetic acid and 1mM DDT). Immunoprecipitates were resuspended in 200µl 25mM pNPP in phosphatase buffer, and incubated for 30 min at 37°C. Reactions were terminated by adding 800µl 1M NaOH, and SHP-1 activity was determined by measuring absorbance at 405nm.

### Cytotoxicity assay

The cytolytic activity of NK cells against target cells was determined with a standard [^35^S]Met release assay. Target cells were labeled with [^35^S]Met (0.2 mCi/ml) for 12 to 16 hours and washed two times, and then 5X10^3^ cells were mixed with NK cells at an effector-to-target ratio of 10:1. Cells were then incubated for 5 hours at 37°C in complete medium. The cells were centrifuged at 200g for 5 min, the supernatant was mixed with scintillation liquid, and radioactive signal was measured with a βcounter (Packard). Spontaneous release of [^35^S]Met from an equal number of target cells was determined by adding 100 ml of complete medium to target cells that were incubated without NK cells. Maximal release was determined by adding 100 ml of 0.1 M NaOH to an equal number of target cells in the absence of NK cells. Finally, the percentage cell lysis caused by the NK cells was calculated using the following equation: % specific lysis = [(sample signal – spontaneous release)/(maximal release – spontaneous release)] × 100.

### DNA Constructs and Mutagenesis

Human SHP-1 wt cDNA was obtained from Addgene. The cDNA of SHP-1 was subcloned into the expression vector pEYFP-N1(Clontech) to obtain the chimeric protein YFP-SHP-1; pECFP-N1 (Clontech) was subcloned into the YFP-SHP-1 expression vector to obtain the chimeric protein YFP-SHP-1-CFP. To avoid localization of SHP-1 to the nucleus, YFP-SHP1-CFP was mutated at its NLS sequence(He *et al*., 2005). Molecular mutants were prepared using the QuikChange II XL site-directed mutagenesis kit (Stratagene). The mCherry plasmid was previously described(Pauker, 2012).

### Cell transfection and FACS analysis

YTS-2DL1 or 721.221/K562 cells were transfected with Nucleofector 2b (Lonza) using Amaxa solution R and protocol X-001. Transiently transfected cells were used after 24-48 hrs. Cells transiently expressing chimeric proteins were selected in hygromycin. Fluorescence analysis and cell sorting were performed using FACSAria or FACSVantage (Becton Dickinson Biosciences).

### Cell stimulation, immunoblotting and immunoprecipitation

First, NK cells (primary or YTS cell line) and 721.221 target cells (either expressing HLA-Cw4, Cw7 or no HLA) were incubated separately on ice for 10 min, at a ratio of 1:1. The cells were mixed, centrifuged and incubated on ice for 15 min. The cell mixture was then transferred to 37°C for the indicated period of time, and subsequently lysed with ice-cold lysis buffer (1% Brij, 1%n-Octyl-b-D-glucoside, 50 mM Tris–HCl, pH 7.6, 150 mM NaCl, 5 mM ethylenediaminetetraacetic acid, 1mM Na_3_VO_4_, and complete protease inhibitor tablets). YTS-2DL1 cells were incubated for 30 min on ice with the indicated concentration of pervanadate, before and at the time of NK-target cell co-culture.

For analysis of whole cell lysates (WCL), 1-5 x 10^5^ cells were used, and for IP experiments 10-15×10^6^ cells were used. Protein A/G plus-Agarose beads (Santa Cruz Biotechnology) were used for IP. Protein samples were resolved with sodium dodecyl sulfate-polyacrylamide gel electrophoresis (SDS-PAGE), transferred to nitrocellulose membrane, and immunoblotted with the appropriate primary antibodies. Immunoreactive proteins were detected with either anti-mouse or anti-rabbit horseradish peroxidase-coupled secondary antibody followed by detection with enhanced chemiluminescence (PerkinElmer).

For WCL samples, the phosphorylation or expression level of proteins was measured by densitometric analysis relative to the GAPDH loading control, using ImageJ. For IP samples, the relative binding or phosphorylation level was measured by densitometric analysis relative to the precipitation control, using ImageJ.

### Measurement of intracellular calcium concentration

First, 0.5-1×10^6^ NK cells were incubated with 5μM Indo-1-acetoxymethylester (Indo-1-AM, Teflabs) and 0.5mM probenecid (MPB) in RPMI 1640 medium at 37°C for 45 min. The cells were washed once, resuspended in RPMI 1640 without phenol red containing 10mM HEPES and 0.5mM probenecid, and maintained at room temperature for 20 min. The cells were incubated at 37°C for 5 min before measurements, then mixed 1:1 with 721 target cells, and the Ca^2+^ influx was measured by spectrofluorimetry using the Synergy 4 Microplate Reader (Bio Tek).

### CD107a degranulation assay

NK cells (3×10^5^) were co-incubated with 6×10^5^ target cells expressing mCherry at 37°C for 2 hours in the presence of 2µM monensin (BioLegend). The cells were centrifuged, incubated with 1:1000 diluted anti-CD107a for 30 minutes on ice and washed twice. Cells were then stained with isotype-specific AlexaFluor-conjugated antibody on ice for 30 minutes. Cells were washed twice, and analyzed by FACS. YTS-2DL1 or pNK-2DL1 cells were distinguished from the target cells based on mCherry expression by the targets.

### Xenograft mice model

NOD-Rag1^null^IL2rg^null^ (RAG) mice were purchased from the Jackson Labs. All mice used were from colonies that were inbred and maintained under SPF conditions at the Bar-Ilan animal house. Housing and breeding of mice and experimental procedures were approved by the Bar-Ilan University Animal Ethics Committee. 6-8 week old female RAG mice were subcutaneously injected, between the shoulders, with 3×10^6^ 721-Cw4 or Cw7 HLA expressing tumor cells, in 0.1 ml of PBS and Matrigel (CORNING) (1:1 ratio). Mice were inspected daily for general well-being, and at the first indication of morbidity (weight loss, lethargy, ruffled fur), or when they reached 8 weeks following inoculation, they were euthanized by CO_2_. Mice tumor diameters were measured daily with a digital caliper. Tumor volumes were calculated according to the formula:

*Tumor volume (mm^3^) = (smallest diameter^2^ × largest diameter) / 2*

When the tumor volumes reached 75mm^3^, mice were intratumorally administered either 3*10^6^ irradiated W.T YTS-2DL1 or SHP-1 deficient NK cells that were untreated or treated with PKC-θ siRNA. Intratumor injections of NK cells were repeated every 3 days for 6 injections in total.

Average growth rates of the tumor from initiation of treatment were calculated according to the formula:

*Average growth rate = (Days of treatment) / (Last day tumor size - Initial tumor size)*

### Statistical analyses

Data calculations of Mean±SE were conducted in Microsoft Excel (v14.7.2), while data were graphed and statistical analysis was performed using GraphPad Prism 9.0.1 (GraphPad Software, Inc., USA). P values were calculated using a two-tailed unpaired T test or One sample t-test. Where >2 conditions were compared, a one-way ANOVA or two way ANOVA with a Tukey post-test was used to calculate P values. Data are depicted as columns with SE. Statistical parameters and biological replicates are reported in the figure legends.

## Data availability

All data generated or analysed during this study are included in the manuscript and supporting file; Source Data files have been provided for Figures 2, 3,5,6,7,8 and supplement figures 1A, 2A and 3A.

## Acknowledgments

The authors thank Danielle Keizer and Dr. Itay Lazar from Bar-Ilan University for their technical assistance. We thank Dr. Jennifer Benichou Israel Cohen for her help with the statistical analysis. This research was partially funded by the Israel Science Foundation.

## Author contributions

MBS, ABS and BS designed the research. ABS, BS, GB, AP, ML, TJ, NJ, OM and JK performed the experiments. ABS, BS, GB, and MBS analyzed the data. MBS and ABS wrote the paper.

## Supplementary figures

**Figure 1-Figure Supplement 1:**
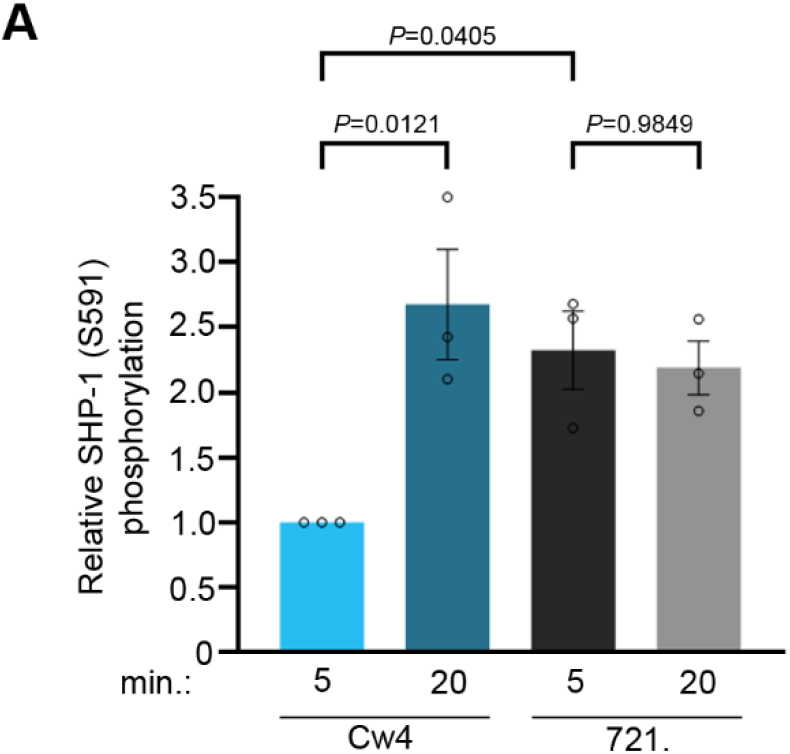
Phosphorylation of SHP-1 S591 is mediated through PKC-θ. **(A)** pNK-2DL1 cells were incubated with target cells as described in Fig. 1C, for two different activation time points, as indicated. pSHP-1 S591 levels were determined as in Fig. 1C. pSHP-1 S591 levels of pNK-2DL1 cells incubated with targets for 5 and 20 minutes are shown on the bar graph. The bar graph shows the average of 4 independent experiments.

**Figure 3-Figure Supplement 1:**
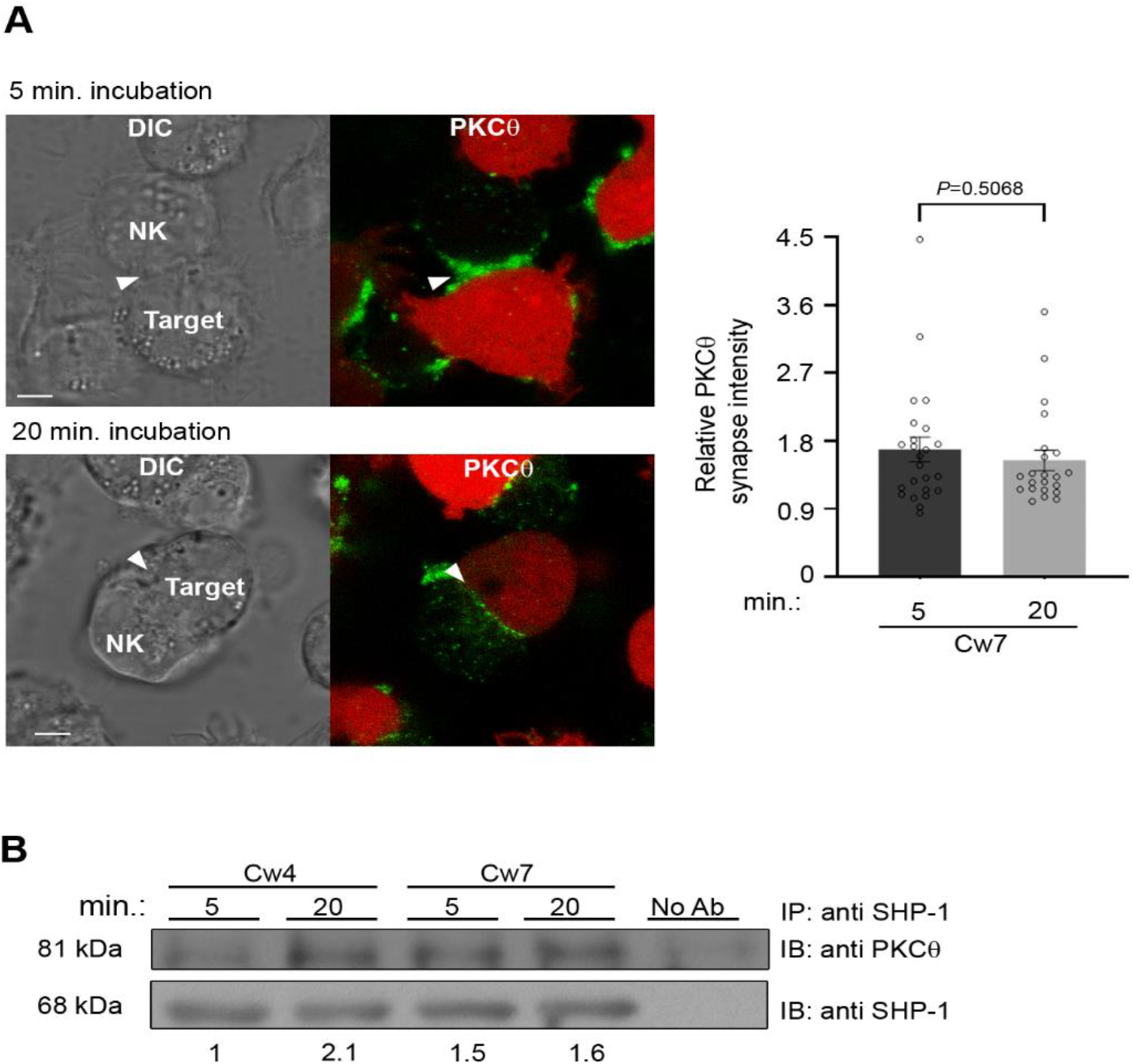
Phosphorylation of SHP-1 is mediated through PKC. **(A)** YTS-2DL1 cells were incubated over slides pre-seeded with 721-Cw7 target cells expressing mCherry. The cells were incubated for 5 or 20 minutes at 37°C to enable conjugate formation, and were fixed. PKC-θ was subsequently labeled with primary goat anti PKC-θ antibody, and secondary anti-goat 488 antibody. Graph summarizing PKC-θ accumulation at the NKIS during each activation time. Analysis was conducted comparing PKC-θ intensity at the NKIS relative to the rest of the NK cell. For Cw7, 5 and 20 minute activation, n=27, and 24 conjugates were analyzed. **(B)** YTS-2DL1 cells were incubated with 721.221 targets for the indicated times. Subsequently, cells were lysed, and lysates were immunoprecipitated with anti-SHP-1 antibody. Lysates were then separated on SDS page and immunoblotted with anti PKC-θ antibody or anti SHP-1 antibody as a loading control. Densitometry was performed relative to YTS-2DL1 sample vs. 721-Cw4 after 5 min activation.

**Figure 4-Figure Supplement 1:**
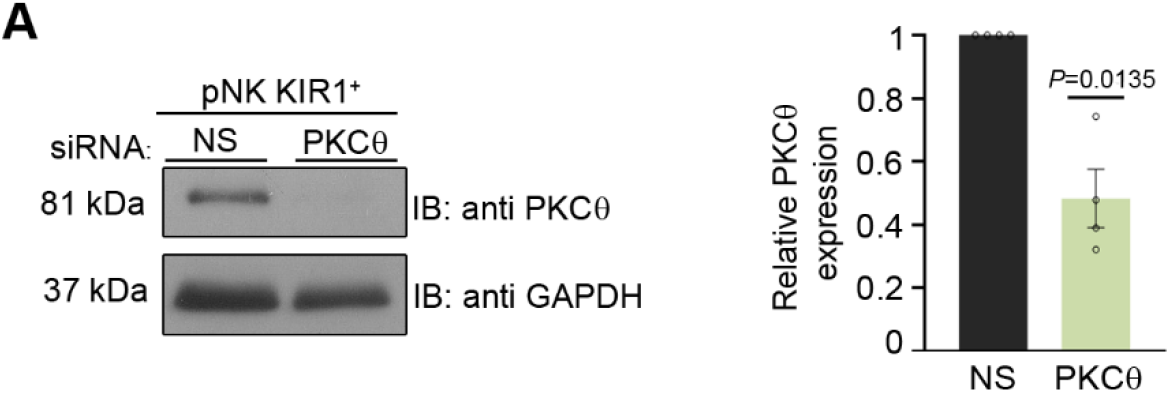
**(A)** Silencing efficiency of PKC-θ. pNK-2DL1 cells were transfected with 250 pmol of PKC-θ siRNA. After 48hrs, prior to incubation with target cells, pNK-2DL1 were counted and lysed. Lysates were separated on SDS page and immunoblotted with anti PKC-θ antibody. PKC-θ expression levels were measured by densitometric analysis, relative to the GAPDH loading control using ImageJ. Samples were normalized according to the pNK-N.S siRNA sample. Bar graph on the right shows the average of data pooled from 3 independent experiments. Data are shown as mean ± SEM. one-way ANOVA with Tukey test (A) or one-sample t test (B, D) were used to calculate p values.

**Figure 5-Figure Supplement 1:**
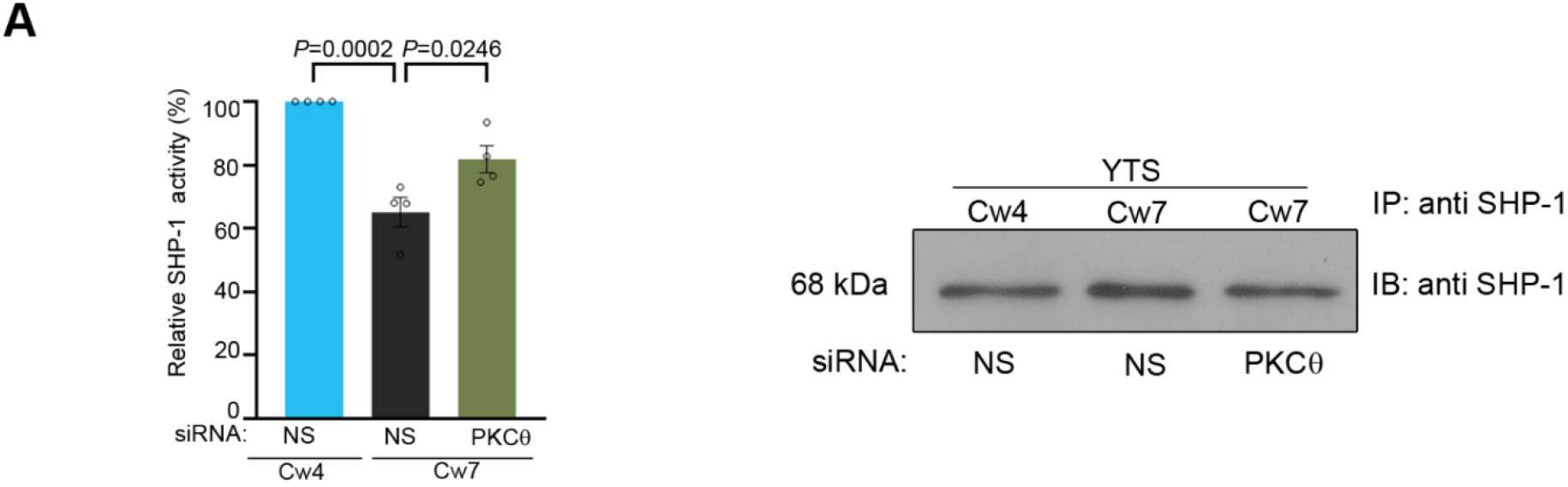
The role of PKC-θ in NK cell activation. **(A)** YTS-2DL1 cells were treated with N.S or PKC-θ siRNA for 48 hrs, and SHP-1 activity was determined. Loading control of SHP-1 precipitated for the PTP assay is shown to the right. The bar graph represents averages of 4 independent experiments.

**Figure 6-Figure Supplement 1:**
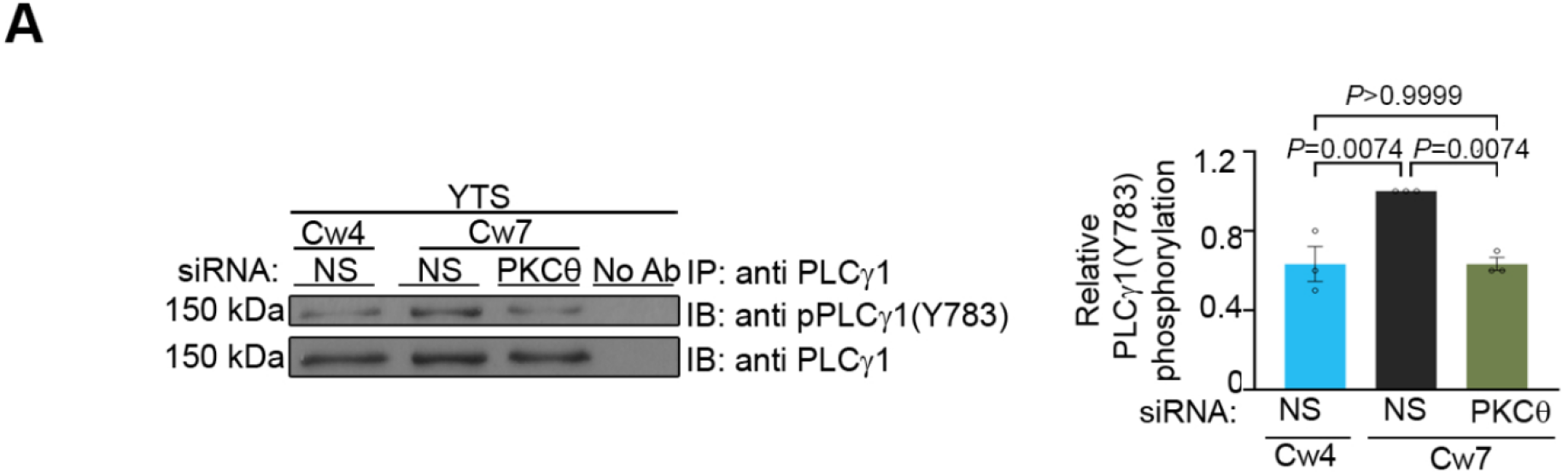
**(A)** YTS-2DL1 cells were transfected with either N.S or PKC-θ siRNA 48 hrs prior to the experiment. Cells were incubated with either 721-Cw4 or Cw7 target cells for 5 minutes at 37°C, and subsequently lysed. Lysates were immunoprecipitated on beads containing PLCγ-1 antibody and immunoblotted for pPLCγ-1(Y783). Densitometric analysis was normalized to PLCγ-1 loading controls, and relative to the YTS-2DL1 N.S siRNA: Cw7 pPLCγ-1(Y783) sample. Bar graph on the right shows the average of data pooled from 3 independent experiments. Data are shown as mean ± SEM. one-way ANOVA with Tukey test (A, B) were used to calculate p values.

**Figure 7-Figure Supplement 1:**
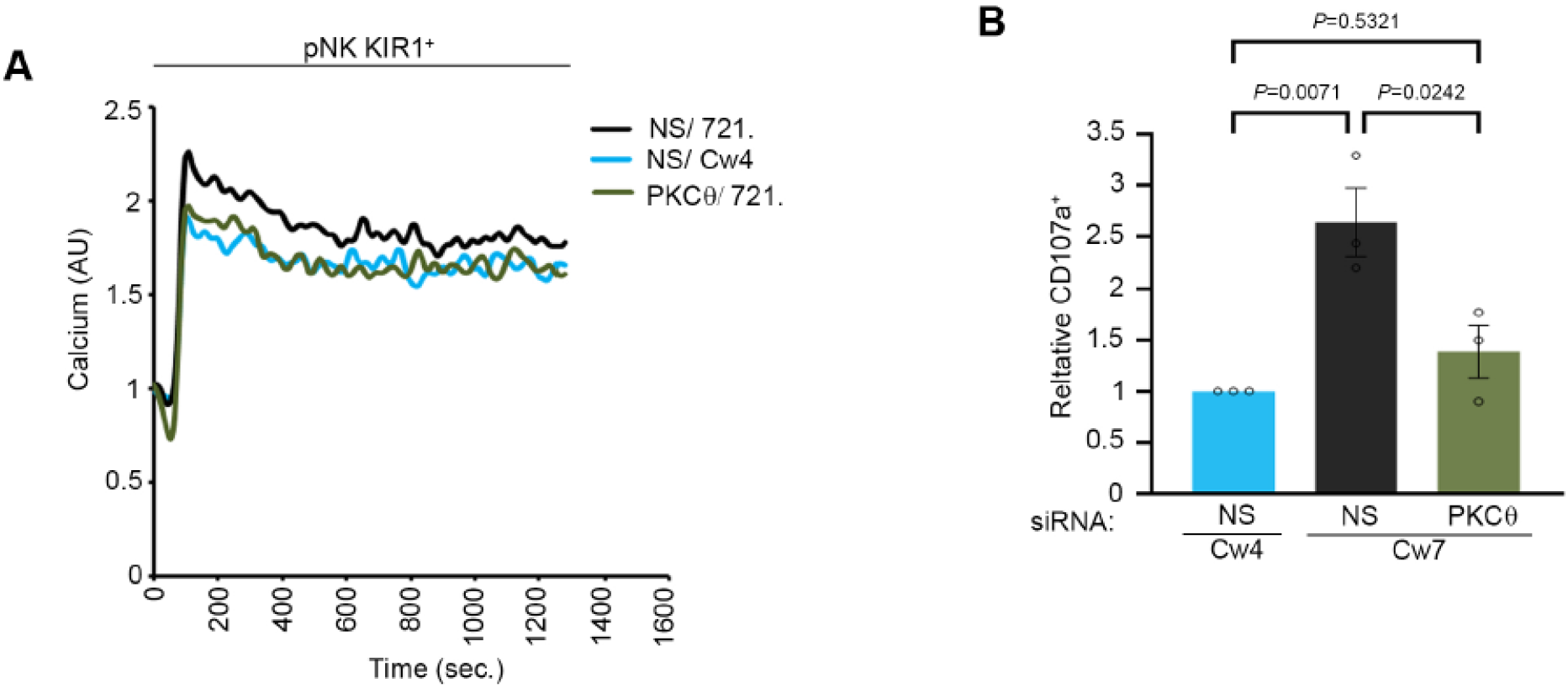
NK cell activation threshold is increased following gene silencing of PKC−θ. **(A)** pNK-2DL1 cells were transfected with specific PKC-θ siRNA or with N.S siRNA, and were used after 48 hrs. pNK-2DL1 cells were loaded with Fluo-3-AM, and analyzed for basal intracellular calcium levels for 1 minute. The NK cells were then mixed with 721-Cw4 or 721-HLA negative target cells, incubated at 37°C and analyzed by spectrofluorometry. **(B)** pNK-2DL1 cells were PKC-θ gene silenced or treated with non-specific siRNA. pNK-2DL1 cells were incubated with mCherry-expressing 721-Cw7 or Cw4 target cells for 2 hours at 37°C and analyzed for degranulation via FACS. Bar graph on the right is normalized to the pNK-2DL1 vs. Cw4 sample, and represents averages of 3 independent experiments. Data are shown as mean ± SEM. one-way ANOVA with Tukey test (A, B) were used to calculate p values.

